# Direct lipid interactions control SARS-CoV-2 M protein conformational dynamics and virus assembly

**DOI:** 10.1101/2024.11.04.620124

**Authors:** Mandira Dutta, Kimberly A. Dolan, Souad Amiar, Elijah J. Bass, Rokaia Sultana, Gregory A. Voth, Stephen G. Brohawn, Robert V. Stahelin

**Affiliations:** Department of Chemistry, The University of Chicago, Chicago, IL 60637; Department of Molecular & Cell Biology, Department of Neuroscience, California Institute for Quantitative Biology (QB3), Biophysics Graduate Program, University of California Berkeley, Berkeley, California 94720, USA; Borch Department of Medicinal Chemistry and Molecular Pharmacology and the Purdue Institute of Inflammation, Immunology and Infectious Disease, Purdue University, West Lafayette, Indiana 47907; Chicago Center for Theoretical Chemistry, Institute for Biophysical Dynamics, and James Franck Institute, The University of Chicago, Chicago, IL 60637

**Author notes:** equal contribution (Mandira Dutta, Kimberly A. Dolan, Souad Amiar). **co-correspondence:**, or.

## Abstract

M is the most abundant structural membrane protein in coronaviruses and is essential for the formation of infectious virus particles. SARS-CoV-2 M adopts two conformations, M_short_ and M_long_, and regulated transition between states is hypothesized to coordinate viral assembly and budding. However, the factors that regulate M conformation and roles for each state are unknown. Here, we discover a direct M-sphingolipid interaction that controls M conformational dynamics and virus assembly. We show M binds Golgi-enriched anionic lipids including ceramide-1-phosphate (C1P). Molecular dynamics simulations show C1P interaction promotes a long to short transition and energetically stabilizes M_short_. Cryo-EM structures show C1P specifically binds M_short_ at a conserved site bridging transmembrane and cytoplasmic regions. Disrupting M_short_-C1P interaction alters M subcellular localization, reduces interaction with Spike and E, and impairs subsequent virus-like particle cell entry. Together, these results show endogenous signaling lipids regulate M structure and support a model in which M_short_ is stabilized in the early endomembrane system to organize other structural proteins prior to viral budding.

## Introduction

Coronaviruses are lipid envelope-encapsulated single-stranded RNA viruses with a genome encoding four structural proteins – spike (S), membrane (M), envelope (E), and nucleocapsid (N). The lipids that comprise the envelope are derived directly from host membranes at the site where viral budding occurs^1^. Viruses often utilize specific lipids to drive every step of the infection cycle – from viral entry^2–5^, to biosynthesis ^6,7^, assembly and egress^8,9^ – and have been shown to modulate the composition of cellular membranes in order to facilitate these processes^10^. Lipids are known to play multiple roles in betacoronavirus infection. Mouse hepatitis virus and SARS-CoV-2 target cholesterol- and ceramide-rich microdomains for viral entry^5^ and depletion of cholesterol reduces S receptor binding in cells^11^. Coronavirus genome amplification and biosynthesis are marked by the formation of double-membrane vesicles enriched in triacylglycerols and cholesterol^6,12^. Coronavirus N proteins, whether free or bound to viral RNA, associate with lipid membranes and specific interactions with anionic lipids may drive localization to sites of viral transcription and assembly^13–15^.

SARS-CoV-2 M is a highly conserved structural membrane protein that facilitates virus or virus-like particle assembly through direct protein-protein interactions with the other structural proteins S, E, and N^16,17^. Coronavirus M proteins adopt distinct conformations, M_short_ and M_long_, in viral envelopes and cryo-EM structures of SARS-CoV-2 M have been captured in each state^16,18,19^. The transition between conformations has been speculated to be important for viral assembly or budding^16,19^. It is hypothesized that interactions between the M protein and specific lipids are critical for these conformational transitions. Recent AFM studies revealed a thinning of membrane thickness near the M protein dimer^20^. Despite the M protein’s critical roles in infectious viral particle assembly and release, and the growing evidence of the critical importance of lipids in these processes, potential roles for specific M-lipid interactions remain unstudied.

Here, we discover M directly interacts with sphingolipids including ceramide-1-phosphate (C1P) in a conformationally selective manner and show that the M-C1P interaction is critical for viral assembly.

## Results

### M binds anionic sphingolipids

Virus assembly and budding often require lipid-protein interactions for assembly site formation and budding and egress of virus particles. To determine if the M protein binds specific lipids, we screened purified M protein for binding to an array of sphingolipids. HA-tagged M protein robustly interacted with C1P as well as dihydro-C1P with appreciable binding to sphingosine-1-phosphate (So1P) and sphinganine-1-phosphate (S1P) (Fig. 1a). To control for construct-specific effects, we also tested a 50:50 mixture of EGFP-M:untagged M and His-SUMO-tagged M protein. Under all of these conditions, M displayed the most robust binding to C1P (Fig. 1a). M did not show appreciable binding to a host of other lipids tested (Supplementary Fig. 1a,b). M did not show appreciable binding to other lipids (Supplementary Fig. 1a,b). To confirm M-C1P binding in a complementary assay, we compared M binding to C1P-coated beads, anionic phosphatidic acid (PA)-coated beads, and control uncoated beads. M protein displayed appreciable binding to C1P-coated beads over control and PA-coated beads (Fig. 1b).

**Figure 1.**
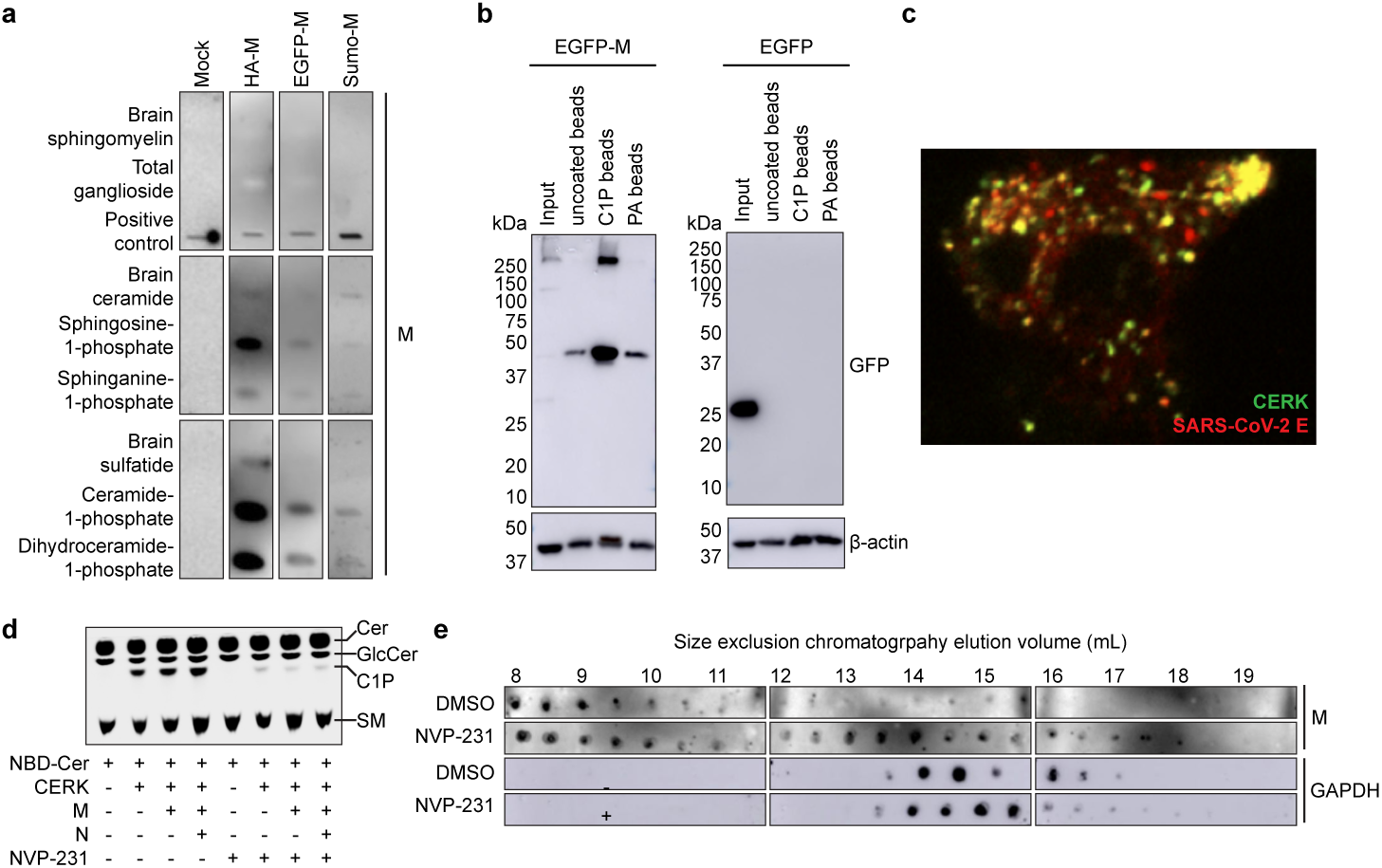
SARS-CoV-2 M protein interacts with C1P. (a) Sphingolipid snooper assay for M-lipid interactions. Western blot of immobilized lipids probed with HA-tagged M, a 50:50 mixture of EGFP-tagged and untagged M, His- and Sumo-tagged M, or a control sample without M protein. All M constructs tested displayed strong binding to C1P and dihydroC1P as well as appreciable binding to anionic So1P and Sa1P. (b) Bead pull down assay of M-lipid interactions. Western blot of input EGFP-M (left) or EGFP (right) expressing cell lysate or eluant from control uncoated, C1P-coated, and PA-coated beads. EGFP-M protein was robustly pulled down by C1P-coated beads. (c) Colocalization of SARS-CoV-2 assembly sites with CERK. HEK293 cells expressing CERK and SARS-CoV-2 structural proteins were imaged. CERK displayed colocalization with fluorescently labeled E protein. (d) Assay for C1P production. Thin layer chromatography of samples from mock-, CERK-, CERK and M-, or CERK, M, and N-transfected HEK293 cells treated with 2.5 µM NBD-ceramide and either DMSO or 100 nM CERK inhibitor NVP-231. CERK increased C1P levels and was inhibited by NVP-231. (e) Assay for M oligomerization in cells. Western blot for M (upper) or GAPDH (lower) in size exclusion chromatography fractionated CERK, M, N, E, and S-expressing HEK293 cell lysate treated with DMSO or 100 nM NVP-231. Later eluting fractions correspond to smaller species. Lower C1P levels altered M protein oligomerization state with an increased proportion of smaller species detected.

Ceramide kinase (CERK) is the major producer of C1P in mammalian cells^21^ and to assess the role of CERK in M assembly, we imaged M and other structural proteins under different conditions (Supplementary Fig. 2). M protein colocalized with the S and E proteins. M protein also colocalized with markers of the endoplasmic reticulum-Golgi intermediate compartment (ERGIC) and Golgi, but not an ER marker (Supplementary Fig. 3a-c), and formed assembly sites similar in vesicle diameter (Supplementary Fig. 3d-f) to authentic SARS-CoV-2^22^. A GFP-CERK construct displayed colocalization with SARS-CoV-2 assembly sites labeling the E protein (Fig. 1c). To assess the role of CERK in M protein’s cellular localization, we expressed CERK, CERK and M, or CERK, M, and N in HEK293 cells and assessed C1P content using thin layer chromatography. C1P production was increased upon CERK expression(Fig. 1d). Next, we treated cells with the CERK inhibitor NVP-231 to confirm reduction in C1P levels, which was similarly lowered under all conditions with CERK expression (Fig. 1d). We then expressed M, S, E, and N with CERK in cells and fractionated lysates using size exclusion chromatography (Figure 1e). Inhibition of CERK resulted in an increase in later-eluting, smaller M oligomers, demonstrating a reduction in higher order oligomerization of M protein when C1P was depleted.

### C1P binds and stabilizes M in the short conformation

We used molecular dynamics (MD) simulations to better understand the nature and consequences of M-C1P interactions. Atomistic models of M were constructed from M_short_ (7VGS) and M_long_ (7VGR) structures embedded in lipid membranes^19^. Membrane composition mimicked the composition of the ERGIC (45% PC, 20% PI, 15% PE, 10% Cholesterol) with either 10% C1P or 10% PS as a control anionic lipid^23,24^. The conformation of M was monitored during the 4 µs simulations by two collective variables that distinguish M_short_ from M_long_: (1) the angle between cytoplasmic C-terminal domains (CTDs) of each subunit and (2) the distance between hinge regions (Fig. 2a).

**Figure 2.**
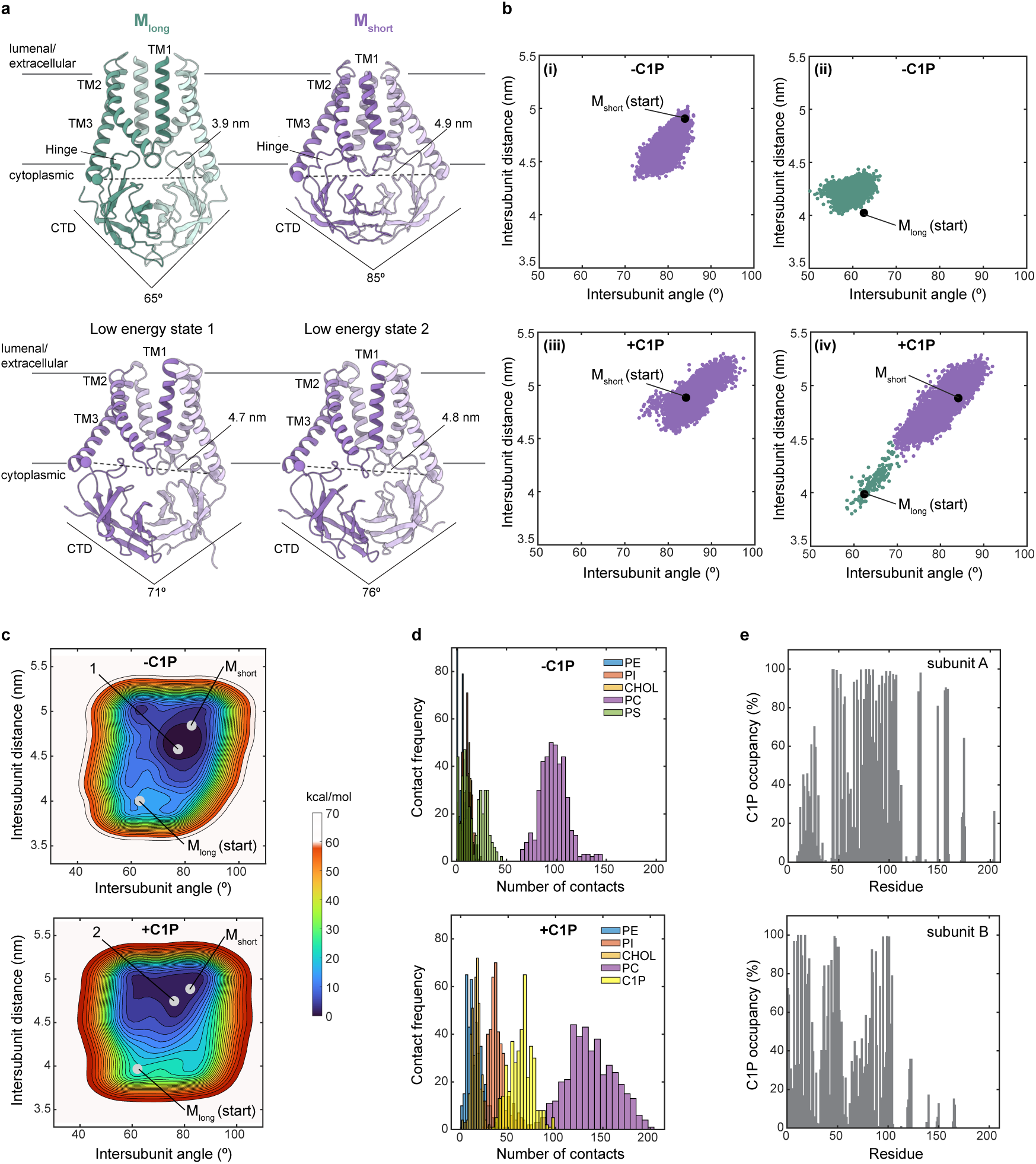
C1P energetically stabilizes M_short_. (a) Structures of M protein viewed from the membrane plane. Collective variables that report on intersubunit distance and angle used to monitor conformational state during simulations are illustrated on the structures. (b) M protein conformational trajectory during molecular dynamics simulations. Intersubunit distance versus intersubunit angle of the structure at each time sample during 4 us simulations is plotted. Simulations were performed without (i,ii) or with C1P (iii, iv) and initiated from M_short_ (i,iii) or M_long_ (i, iv). Conformational switching was only observed from M_long_ to M_short_ in the presence (of C1P. (c) Free-energy landscape calculated from simulations without (above) or with C1P (below). Position of M_short_, M_long_, and a low energy state from each condition is indicated. (d) M-lipid contact frequency from simulations without (above) and with C1P (below). (e) M-C1P percentage occupancy by residue in subunit A (above) and subunit B (below).

Strikingly, MD simulations initiated from M_long_ in the presence of C1P showed rapid conformational switching to M_short_-like conformations, which remained stable for the duration of the simulation (Fig. 2b). Conformational switching was dependent on C1P, as simulations initiated from M_long_ in the absence of C1P did not exhibit a transition to the short state. Notably, switching was only observed from M_long_ to M_short_; M_short_ in the presence or absence of C1P remained in M_short_-like conformations throughout the simulations (Fig. 2b). This is consistent with the stability of M_short_ we observed in prior simulations in the absence of C1P^18^.

These results suggest that C1P stabilizes M_short_. We computed the conformational free energy landscape using well-tempered (WT) metadynamics^25^, employing the same collective variables that utilize the intersubunit CTD angle and hinge distance as the pertinent coordinates (Fig. 2c). We reach two conclusions from this analysis. First, M_short_ is a lower free energy state than M_long_, regardless of lipid composition. Minimum energy states in the absence (indicated by “1”) and presence of C1P (indicated by “2”) are similarly M_short_-like (Fig. 2a). Second, C1P markedly stabilizes M_short_. We calculate the magnitude of stabilization to be –6.99±1.00 kcal/mol (defined as ΔΔ*G* = Δ*G_C_*_1*P*_ − Δ*G_C_*_1*P*_ = (-18.07±0.62)-(-11.08±0.79)} kcal/mol = -6.99±1.00 kcal/mol). Stabilization is primarily due to a deeper free energy well around M_short_-like conformations.

C1P stabilization could be explained by direct binding to M_short_. We observed more CTD-lipid headgroup contacts in MD simulations with C1P, with enrichment of C1P and PI contacts (Fig. 2d). C1P and PI lipids were localized around M_short_ in visualizations of lipid position (Supplementary Fig. 4a). Analysis of membrane structure during the last 2 μs of the simulations with C1P shows M subtly thins and curves the membrane, particularly around the CTD-inner leaflet interface (Supplementary Fig. 4b-e). To identify candidate residues involved in M-C1P interactions, we calculated residence time (Supplementary Fig. 4f) and percentage occupancy (Fig. 2e) of C1P around M by amino acid. A continuous patch of residues at the TM-CTD interface showed both high C1P occupancy (>80%) and among the longest C1P residence times. The patch includes residues at the bottom of TM1 (R44, F45, and I48), bottom of TM3, hinge (A104, R107, S108, M109, and W110), proximal CTD (L129 and R131), and nearby charged residues in the CTD (H148, H155, and R158).

We next sought direct experimental structural insight into the molecular basis of M-C1P interaction. We prepared complexes of M with CTD-binding Fabs that stabilize either M_short_ or M_long_ conformations in the presence of 100 µM C1P. To facilitate comparisons to previously reported apo M structures, buffer and detergent conditions were otherwise identical. We determined cryo-EM structures of M_short_ and M_long_ in the presence of C1P to 3.0 Å overall resolution (Fig. 3a, Supplementary Fig. 5,6, Table 1). The M_short_-C1P map shows lipid-like densities not visible in apo-M_short_ maps. One density feature per subunit at the inner leaflet can be unambiguously modeled as C1P with well-defined phosphomonoester headgroup and acyl chains (Fig. 3a). C1P binds to a site formed by the bottom of TM1-3, hinge region, and membrane-proximal aspect of the CTD, corroborating our molecular dynamics simulations (Fig. 3b). The C1P head group fits into an electropositive pocket and forms polar interactions with R44, W110, and R131 (Fig. 3c). C1P acyl chains pack into a cleft between TM2 and TM3, forming hydrophobic interactions with F45, I48, L51, I52, W55, M109, W110, and P114 on TM2 (Fig. 3b,d). Density corresponding to the inner acyl chain is weakly visible in the apo M_short_ map, suggesting partial occupancy by co-purified lipids or detergents in the absence of C1P.

**Figure 3.**
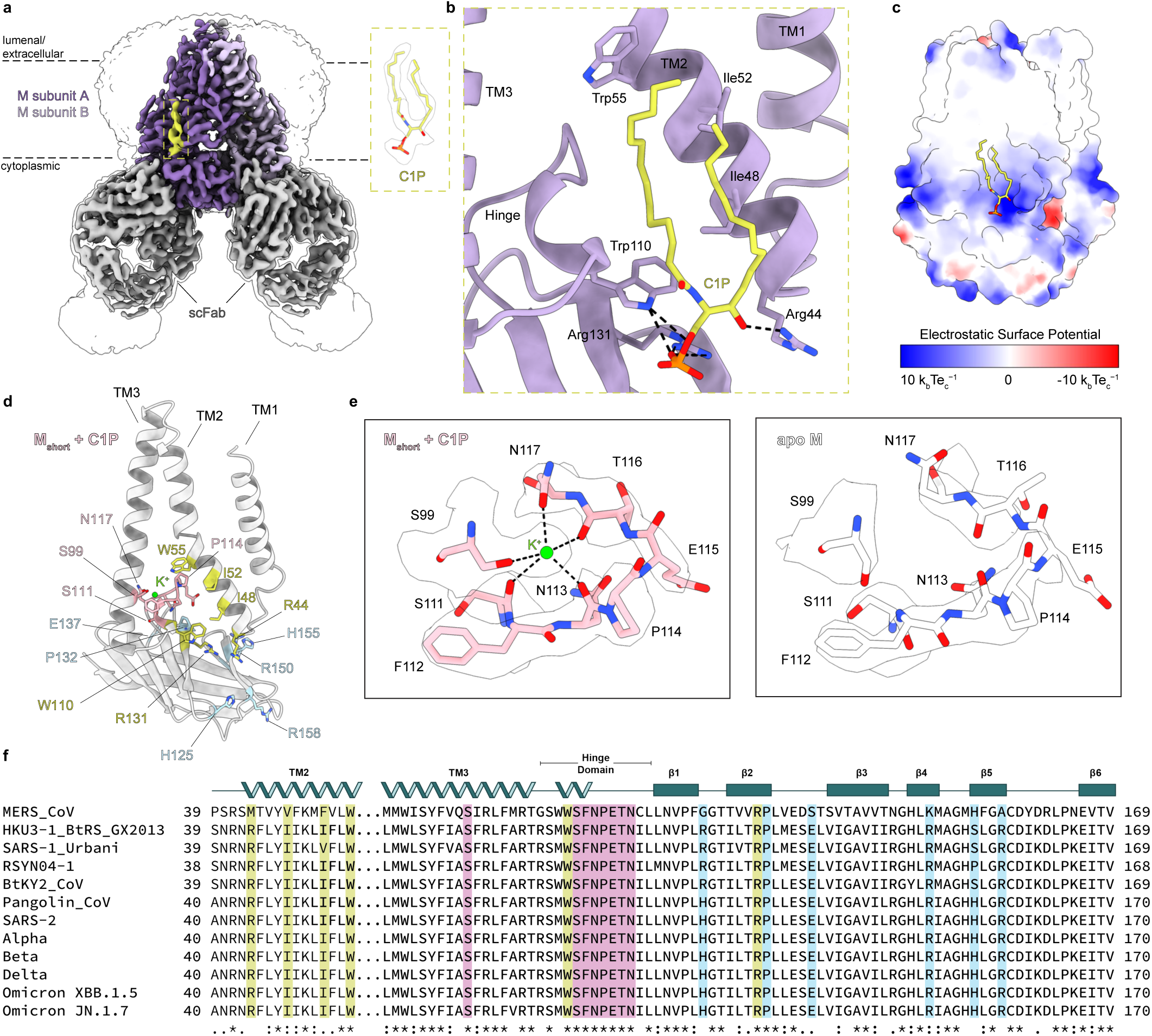
Cryo-EM structure of M_short_ bound to C1P. (a) 3.0 Å resolution cryo-EM map of SARS-CoV-2 M in the short conformation viewed from the membrane. M subunits are purple, short conformation-selective Fabs (scFab) are gray, and C1P is yellow. Unsharpened map is shown transparent at low contour. (b) Zoomed in view of C1P binding site in one subunit. (c) Electrostatic surface view of M with C1P bound. (d) Single M_short_-C1P subunit with C1P binding residues yellow, K^+^ binding residues pink, and residues displaying associated conformational changes blue. (e) Zoomed in view of K^+^ binding site in M_short_-C1P (left) or apo M_short_ (right). Cryo-EM density is shown as a transparent surface. (f) Sequence alignment of M across coronaviruses with residues colored as in (d).

**Table 1.** Cryo-EM data collection, refinement, and validation statistics.

|  | M:scFab + C1P | M:lcFab + C1P |
| --- | --- | --- |
| PDB ID | 9CTU | 9CTW |
| EMDB ID | 45919 | 45921 |
| EMPIAR ID | xxx | xxx |
| **Data collection** |  |  |
| Total movies | 6,741 | 8,432 |
| Magnification | 105,000x |  |
| Voltage (kV) | 300 |  |
| Electron exposure (e-/Å<sup>2</sup>) | 50 |  |
| Defocus range (um) | -0.5 to -1.8 |  |
| Pixel size (Å<sup>2</sup>) | 0.4055 (super res) | 0.848 |
| **Processing** |  |  |
| Initial particle images (no.) | 351,588 | 561,632 |
| Final particle images (no.) | 44,736 | 190,251 |
| Map resolution Masked (Å, FSC = 0.143) | 3.03 | 3.01 |
| Symmetry imposed | C2 | C2 |
| **Refinement** |  |  |
| Model resolution (Å, FSC = 0.143 / 0.5) | 3.0/3.4 | 2.9/3.1 |
| Map-sharpening B factor (Å<sup>2</sup>) | -108.4 | -119.0 |
| **Composition** |  |  |
| Number of atoms | 9714 | 9754 |
| Number of protein residues | 1240 | 1256 |
| Number of ligands | 4 | 0 |
| **RMS deviations** |  |  |
| Bond lengths (Å) | 0.003 | 0.003 |
| Bond angles (Å) | 0.532 | 0.510 |
| **Validation** |  |  |
| MolProbity score | 1.63 | 1.25 |
| Clashscore | 9.58 | 2.07 |
| **Ramachandran plot** |  |  |
| Favored (%) | 97.39 | 96.06 |
| Allowed (%) | 2.61 | 3.78 |
| Disallowed (%) | 0 | 0.16 |
| Rotamer outliers (%) | 1.03 | 0.74 |
| **Mean B factor (Å<sup>2</sup>)** |  |  |
| Protein | 95.42 | 55.62 |
| Ligand | 73.27 | - |

In contrast, we see no evidence for specific C1P binding to M_long_ (Supplementary Fig. 7a). M_long_+C1P and apo-M_long_ structures are nearly indistinguishable (0.79 Å overall r.m.s.d.). Densities consistent with detergent or lipid acyl chains around the inner leaflet are similar in M_long_+C1P and apo-M_long_ maps. The absence of C1P binding to M_long_ is explained by reorganization of the M_short_-C1P binding site. In M_long_, the two halves of the C1P binding pocket (TM1/2 and TM3/hinge/CTD) spread apart, separating key residues R44 and R131 by > 7 Å (Supplementary Fig. 7b).

Comparing M_short_-C1P and apo M_short_ structures reveals four consequences of C1P binding. First, C1P binding compacts M, particularly in the TM region with intersubunit TM2 and TM3 distances 1.5-3.5 Å shorter (Supplementary Fig. 8a). Second, C1P binding stabilizes M_short_. M_short_-C1P has more clearly resolved flexible regions (including in the hinge and TM linkers) and lower, less variable refined B-factors than apo M_short_ (Supplementary Fig. 8c), despite similar particle numbers, processing routines, and overall resolutions. Third, C1P binding reveals an ion binding site bridging the hinge and bottom of TM3 (Fig. 3d,e). Electronegative groups, including S99 and N117 side chain hydroxyls and S111, N113, and T116 backbone carbonyls, surround a strong density feature not visible in the apo-M_short_ map. We model a K^+^ in this site supported by buffer conditions (150 mM KCl), coordination geometry, and multiple validation tools. Fourth, a series of conformational changes propagate from the C1P binding site and alter the CTD surface character. Hinge residue W110 is shifted towards the CTD upon C1P binding, necessitating a shift in neighboring P132 on the β2 strand (Supplementary Fig. 8b). Charged residues including H125 on β1, R158 on β5, and E137, R150 and H155 at the intersubunit CTD interface change rotameric state, altering the electrostatic character of the CTD surface and potentially influencing M interactions with other structural proteins during virus assembly (Supplementary Fig. 8d). We conclude C1P specifically binds and stabilizes M_short_.

Notably, residues involved in C1P-binding and associated conformational changes have been recently implicated in the action of an anti-coronavirus small molecule. JNJ-9676 binds M_short_ (but not M_long_) in a pocket above the hinge between TM2 and TM3, overlapping the potassium ion binding site identified here^26^. JNJ-9676 is reported to prevent virus assembly by trapping M_short_, suggesting an M_short_-M_long_ transition is critical to the viral life cycle. Among JNJ-9676 escape mutants identified in M, two (P132S, S99A) are not readily explained by steric or chemical perturbation of the drug binding site. We reasoned these mutations could disrupt drug binding by promoting M_long_ and that other residues implicated in C1P binding would have similar effects. We used AlphaFold3 to test this hypothesis. Wild-type M was predicted to adopt an M_short_-like conformation (Supplementary Fig. 9), consistent with our structural and MD results. In contrast, mutants at sites implicated in C1P binding or associated conformational changes (H155E, E137R, R150D, W110R, P132W, and S99D) are predicted to adopt increasingly M_long_-like conformations (Supplementary Fig. 9). Among these structures, M_short_-C1P is the most compacted in an MD trajectory from short to long conformations.

### M-C1P binding is critical for virus assembly of the spike protein

Finally, we asked what role C1P binding has on M function in virus assembly. We prepared several mutations of M and assessed their functional consequences. A cationic patch in the M CTD implicated in MD and structural experiments in C1P interaction (residues 146-RGHLRIAGHH-155) was mutated to replace positively charged arginine and histidine residues with alanine. This mutation resulted in loss of M binding to C1P assessed with lipid-coated bead pull down and is referred to as M^ΔC1P^ (Fig. 4a). A second cationic rich region was also mutated (198-RYRIGNYK-205) as it was previously shown to be a trans Golgi retention signal for MERS M protein (M^ΔTGN^)^27^. Similar to the previous report, mutation of the trans Golgi retention signal disrupted M localization (Supplementary Fig. 10) and colocalization with S and E. In a similar fashion, M^ΔC1P^ disrupted M localization (Fig. 4a,b) and colocalization with S and E (Figure 4b-d).

**Figure 4.**
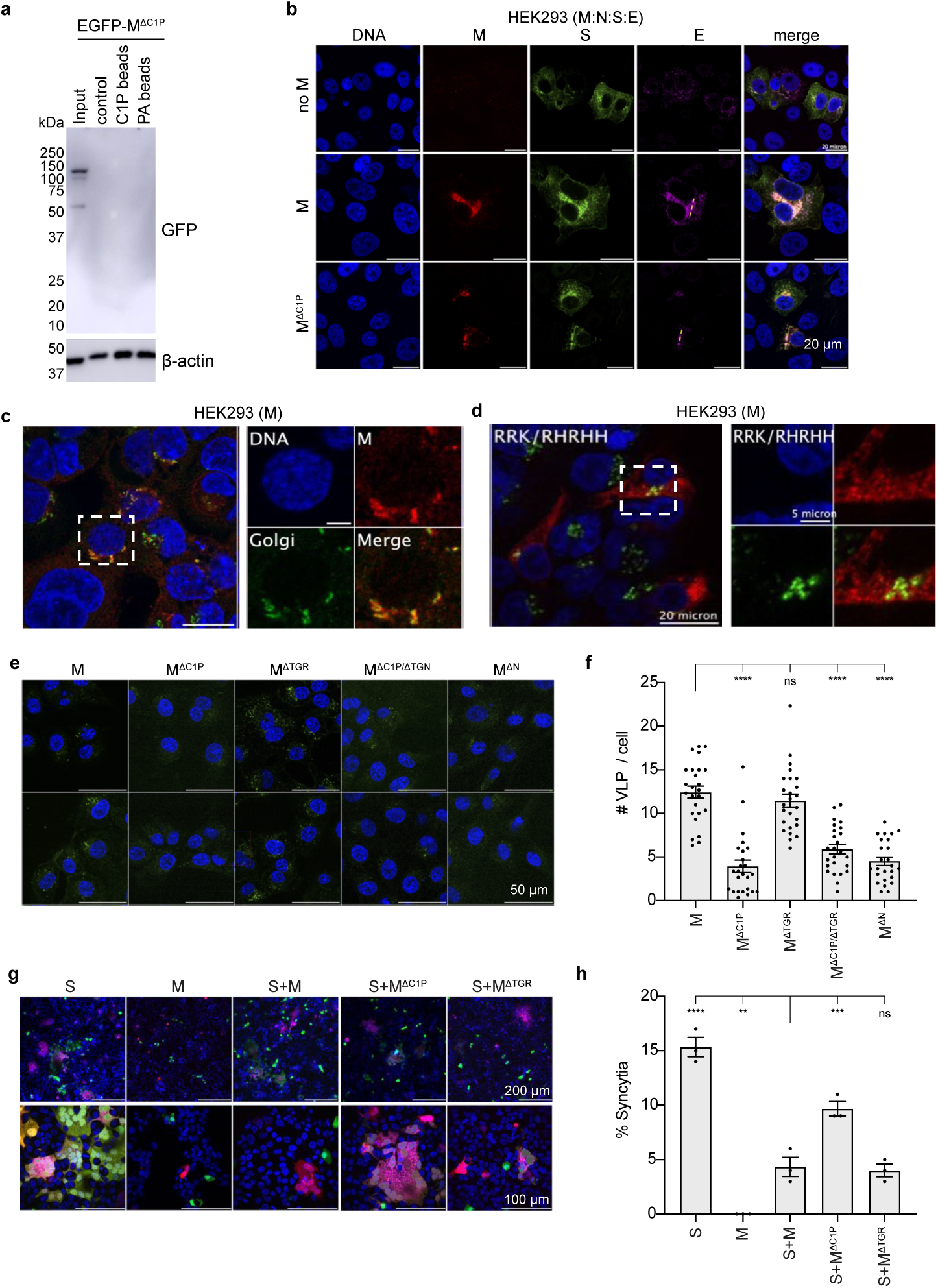
Functional assessment of M protein mutations. (a) Bead pull down assay of M^ΔC1P^ mutant-lipid interactions. Western blot of input EGFP-M^ΔC1P^ expressing cell lysate and eluant from uncoated, C1P-coated, or PA-coated beads. (b) M^ΔC1P^ disrupts colocalization with S observed with wild-type M. (c,d) M^ΔC1P^ disrupts Golgi localization of wild-type M. (e,f) Virus-like particle (VLP) entry assay. M^ΔC1P^, M^ΔC1P/ΔTGN^, and M^ΔN^ (which reduces M:N-interaction) derived VLPs show reduced cell entry compared to VLPs formed with wild-type M or M^ΔC1P^. (g,h) Cell syncytia formation assay. Cells expressing S alone, but not S and M, form robust syncytia. In contrast, co-expression with M^ΔC1P^ significantly increased syncytia formation. Data are mean ± sem with differences assessed with a one-way Annova with Dunnet correction for multiple comparisons (****p<0.0001, ***p<0.001, **p<0.01).

To test the hypothesis that the M-C1P interaction is an important factor for proper S incorporation into VLPs, we assessed VLP entry for WT and mutant M containing VLPs. WT M containing VLPs were able to undergo cell entry (quantified by number of GFP-positive puncta counted per cell) as were VLPs containing M^ΔTGN^ (Fig. 4e,f). In contrast, VLPs derived from M mutations that displayed altered S/M colocalization and reduced C1P binding (M^ΔC1P^) had a significant reduction in cell entry (Fig. 4e,f). This underscores the importance of M-C1P interaction in proper S localization and incorporation into VLPs. To further confirm changes in S interaction and localization, we employed a syncytia formation assay which has been used to assess the trafficking of S to the plasma membrane of cells ^28–30^. S expression alone resulted in robust syncytia formation that was significantly reduced by co-expression with M or M^ΔTGN^ (Fig. 4g-h). In contrast, co-expression of M^ΔC1P^ with S resulted in increased syncytia formation consistent with disrupted S interaction(Fig. 4g-h).

## Discussion

Together, the data presented here support a model for lipid-regulated coronavirus assembly through specific binding and conformational control of SARS-CoV-2 M protein (Fig. 5). We propose C1P binding enriches M_short_ in ERGIC and Golgi membranes to facilitate its organization with other structural proteins S, E, and N in early steps of virus assembly. Transition to M_long_ promotes higher order assembly and virus release and involves loss of C1P through dilution and/or additional promoting factors such as structural protein binding. Several lines of evidence support this model. M is in dynamic equilibrium M_short_ and M_long_^16,19^, with M_short_ energetically favored (Fig. 2)^18^. Specific binding of C1P further stabilizes and compacts M_short_ (Figs. 1-3). C1P is synthesized and enriched in the ERGIC and Golgi membranes ^31^, where M organizes the other structural proteins S, E, and N ^32^ (Fig.1). Depleting C1P results in mislocalization and higher order assembly of M (Fig. 1), while mutations that disrupt M-C1P interaction negatively impact structural protein binding and VLP entry (Fig. 4).

**Figure 5.**
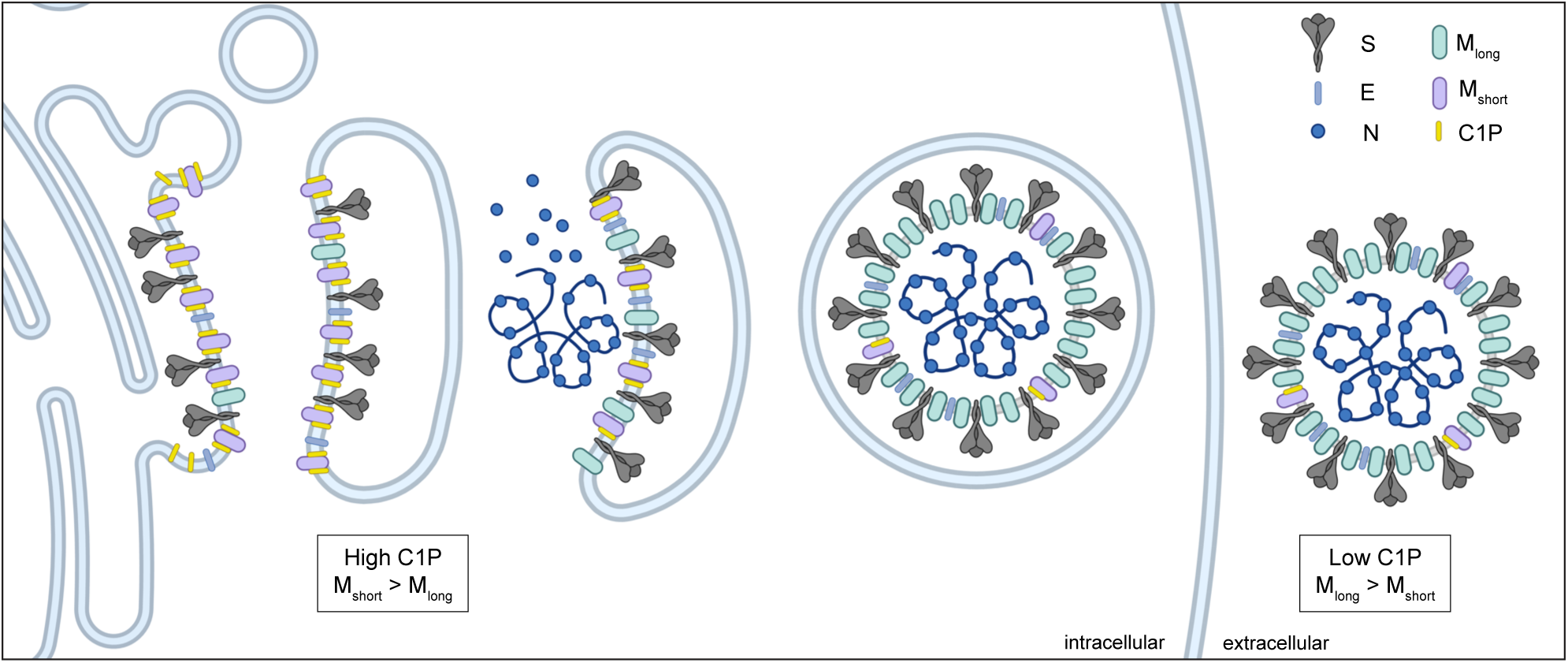
Model for lipid control of M conformational dynamics and coronavirus. assembly. M protein adopts M_short_ and M_long_ conformations. Ceramide kinase localization to ERGIC membranes results in high C1P concentrations. C1P binds and stabilizes M_short_ in the ERGIC to facilitate recruitment of E and S. Decreasing C1P along the secretory pathway and other potential factors trigger a conformational switch to M_long_ that promotes higher membrane curvature and oligomerization necessary for virus assembly and budding.

This model provides insight into the mechanism of action of the recently reported small molecule antiviral JNJ-9676, the first that directly targets M protein^26^. JNJ-9676 mimics the effect of C1P, specifically binding to an M_short_-like conformation and disfavoring transition to M_long_. Unlike M_short_-C1P, M_short_-JNJ-9676 is inhibited from transitioning to M_long_, either because JNJ-9676 is not diluted during progression through the endomembrane systems or because the complex is insensitive to additional unknown M_long_-promoting factors. As a result, M is trapped in an early short-like conformation, preventing formation of mature virus particles. One prediction of this model is that additional M_short_-stabilizing molecules, perhaps additionally co-opting the C1P binding site, are likely to similarly prevent virus formation. A corollary is that M_long_ stabilizing small molecules or antibodies, including those potentially patient-derived^33–37^, are predicted to inhibit virus disassembly and successful infection.

Lipid enveloped viruses hijack host membranes to form sites of virus assembly and give rise to new virus particles. Herein we demonstrate that SARS-CoV-2 M makes direct interactions with host cell C1P, which is critical for conformational stability of M_short_ and correct assembly of S. This illustrates SARS-CoV-2 uses specific host lipid interactions to mediate formation of virus particles akin to other virus families (e.g., filoviruses, retroviruses, paramyxoviruses, flaviviruses, etc. ^38–45^. A number of studies have demonstrated increased cellular levels of ceramide during SARS-CoV-2 infection ^46,47^ where ceramide was shown to help facilitate viral entry ^48,49^. Further, drugs that disrupted ceramide synthesis (e.g., amitriptyline) reduced SARS-CoV-2 entry into cells ^48,50^ and reduced mortality in patients ^51^. While the details of C1P in this process have not been explored in great detail, ceramide is the direct substrate of C1P generation and increases in C1P have been shown in the sera of SARS-CoV-2 infected mice and in infected Vero E6 cells ^52^. Thus, more detailed studies examining ceramide/C1P ratio changes in SARS-CoV-2 infection and cellular models used herein are clearly warranted.

## Figure legends

**Supplementary Figure 1.**
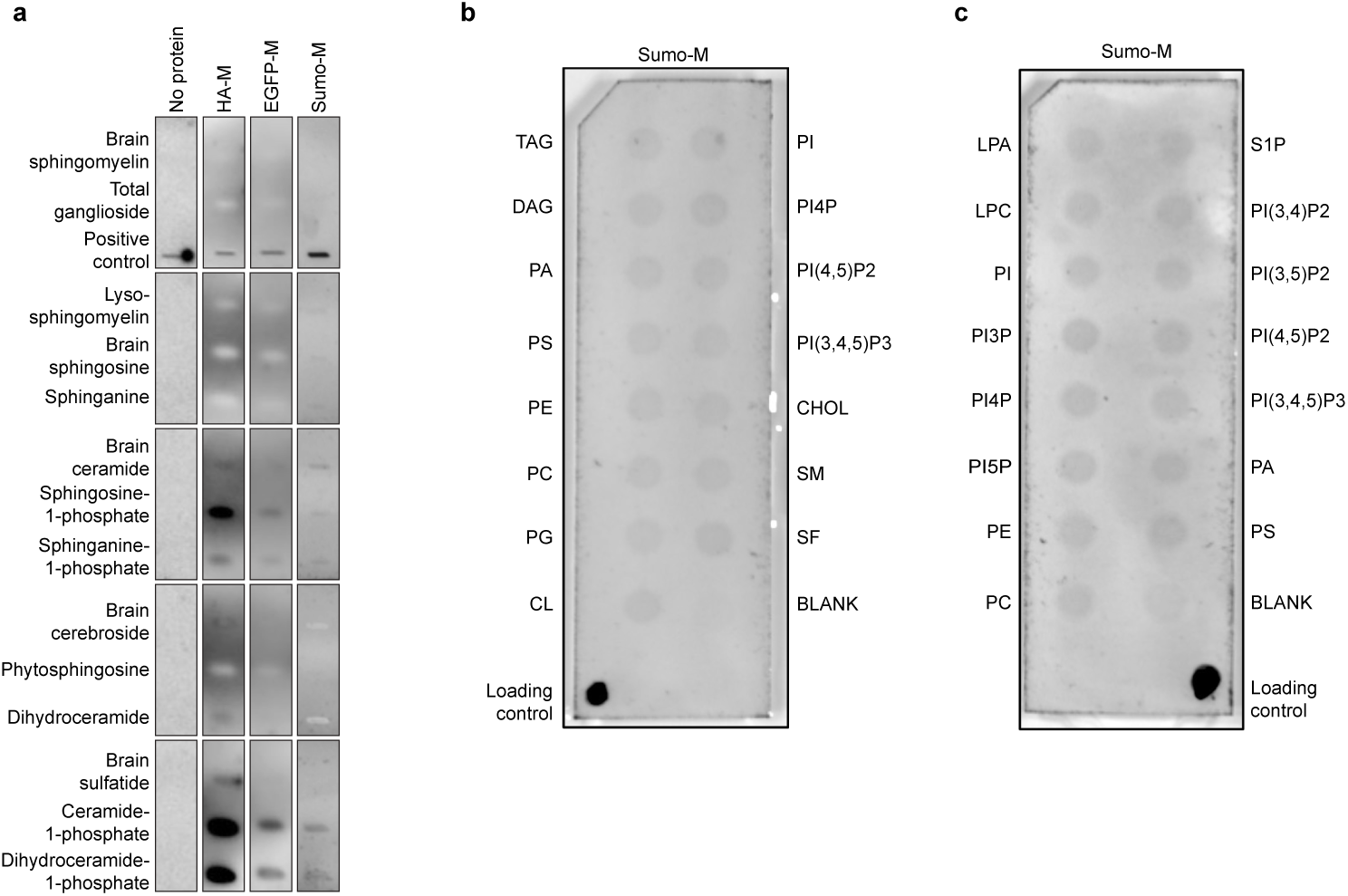
M protein-lipid interactions. (a) Sphingolipid, (b) lipid, and (c) phosphatidyl inositide snooper assays for M-lipid interactions. (a) Western blots of immobilized lipids probed with HA-tagged M, a 50:50 mixture of EGFP-tagged:untagged M, His- and Sumo-tagged M, or a control without M protein. All M constructs tested displayed strong binding to C1P and dihydroC1P as well as appreciable binding to anionic So1P and Sa1P.Lipid and (b,c) Western blots of immobilized lipids probed with His- and Sumo-tagged M. No appreciable binding is observed. All westerns were performed with anti-M antibody and secondary conjugated to HRP. Portions of (a) are reproduced from Fig. 1a.

**Supplementary Figure 2.**
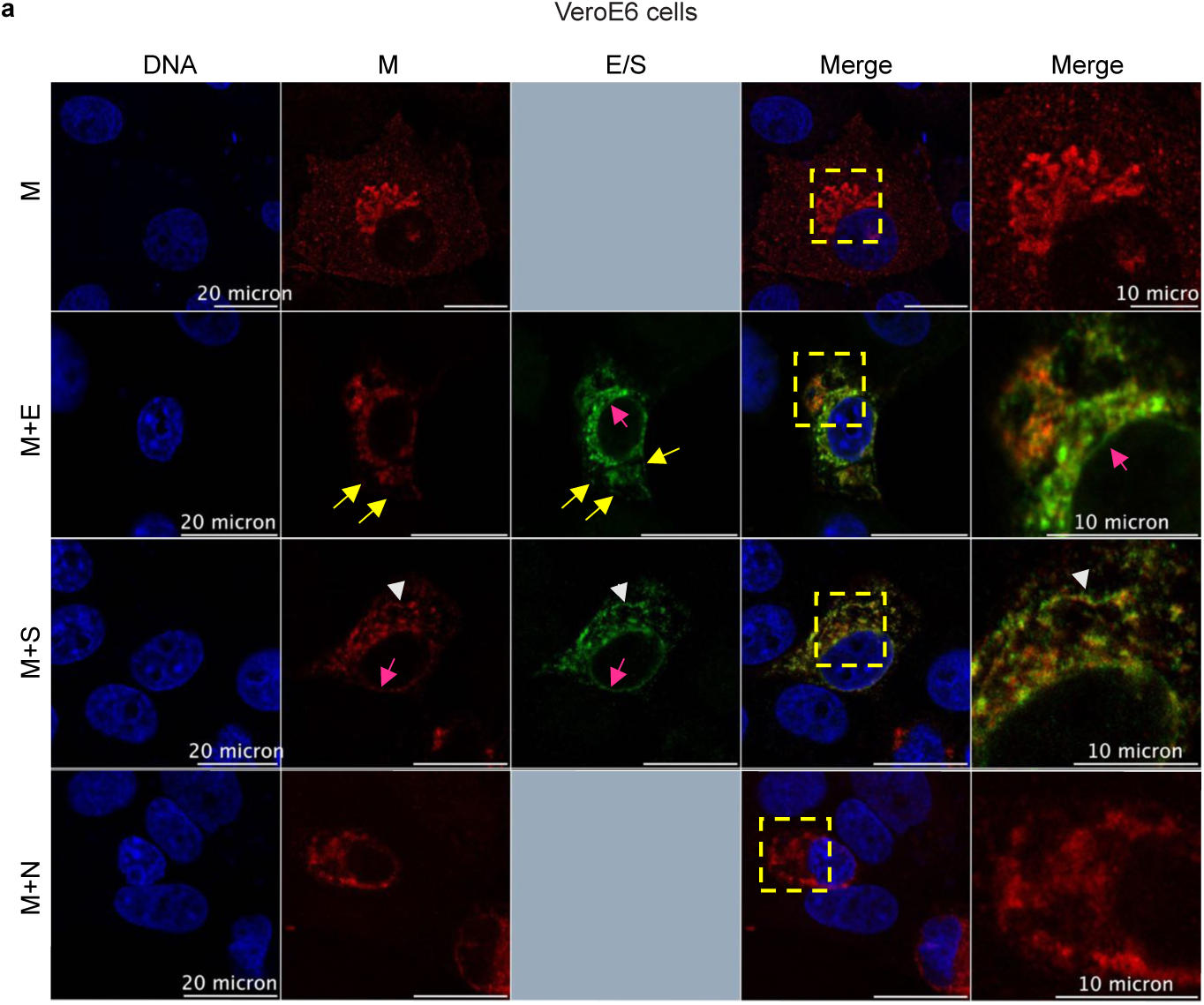
SARS-CoV-2 M colocalizes with S and E proteins. Vero E6 cells were used to assess M protein localization when co-expressed with E, S, or N. The insets and arrows show regions of colocalization.

**Supplementary Figure 3.**
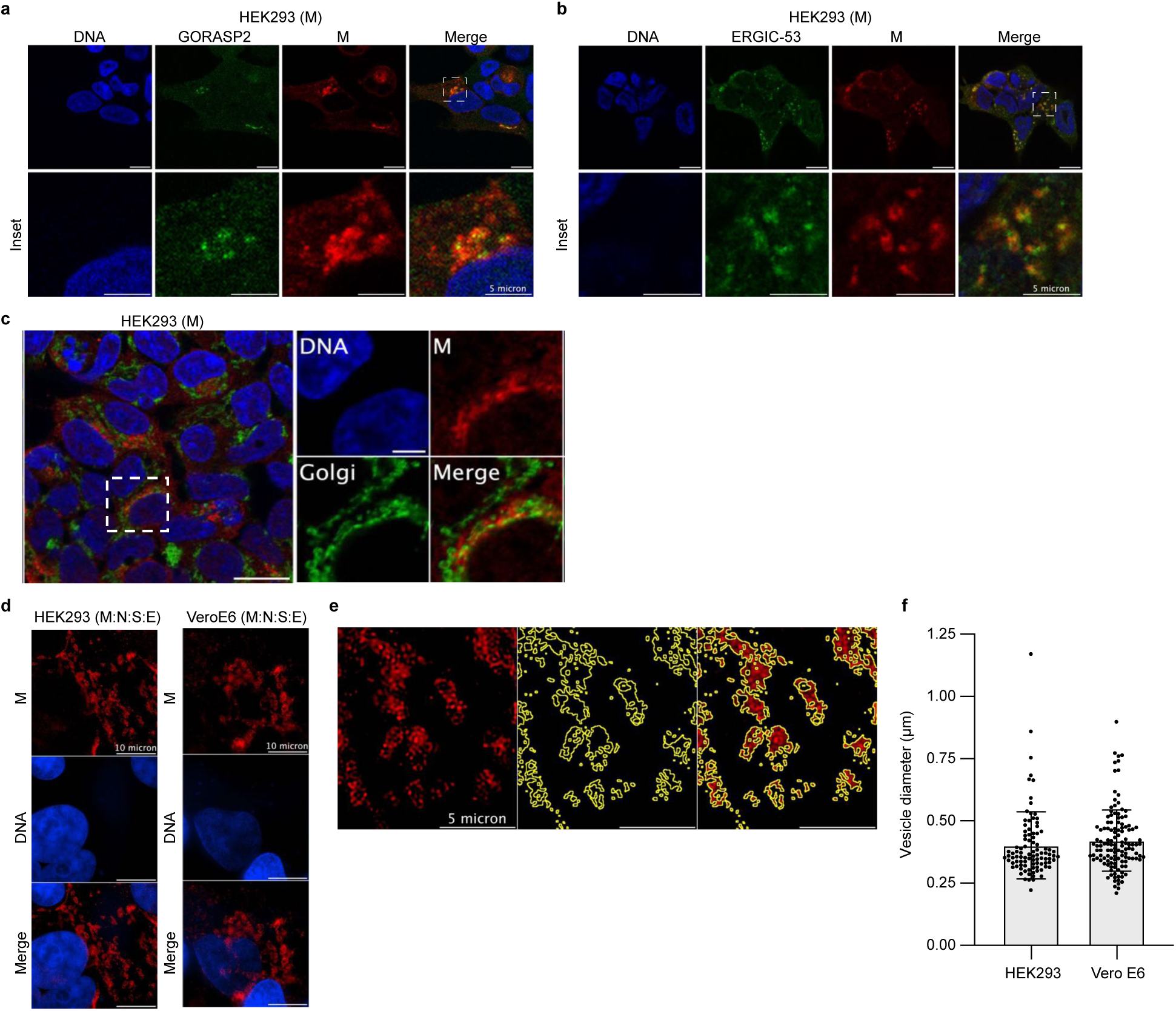
SARS-CoV-2 M colocalizes with Golgi and ERGIC markers and SARS-CoV-2 structural proteins co-expressed in mammalian cells exhibit similar vesicle morphology to SARS-CoV-2 infected cells. (a) WT M protein (red) expressed in HEK293 cells colocalized with the Golgi GORASP2 marker (green). (b) WT M protein (red) expressed in HEK293 cells colocalized with the ERGIC-53 marker (green). (c) WT M protein (red) expressed in HEK293 cells did not appreciably colocalize a ER marker (green) (c) SARS-CoV-2 structural proteins (S, M, Flag-E, and N) were co-expressed in HEK293 or Vero E6 cells and imaged with structure illuminated microscopy using a Nikon N-STORM/N-SIM/TIRF imaging system. Immunofluorescence with labelled anti-Flag antibody for detection of the Flag-E protein (red). (e) A zoomed in portion of (d) is shown to demonstrate how vesicle diameter was quantified. (f) Quantification of vesicle size was performed using ImageJ. Similar average vesicle diameters illuminated with E-Flag antibody were found in HEK293 and Vero E6 cells.

**Supplementary Figure 4.**
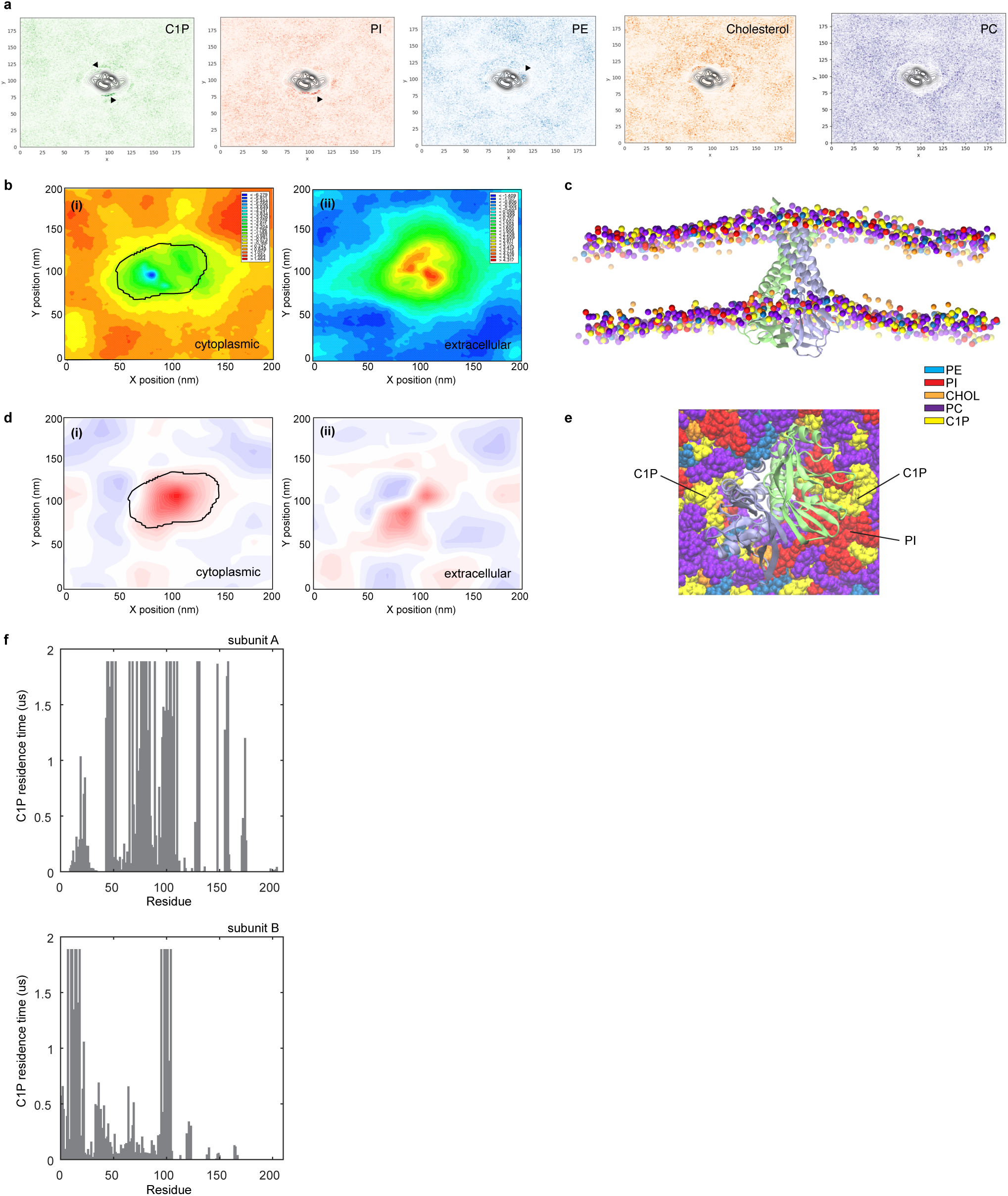
Lipid density map, membrane response, and M-C1P residence time from molecular dynamics simulations. Density for (a) C1P, POPI, DYPE, cholesterol, and PC in the simulation box is illustrated with darker color indicating higher density. Arrowheads indicate higher lipid accumulation. The central region corresponds to M protein. (b) Z-coordinates of lipid head groups averaged over the last 2 μs of trajectories and mapped into the XY plane for outer (i) and inner (ii) leaflets. The black circle in the top leaflet highlights the region of membrane thinning. (b) Side view of the simulated M_short_ box. Membrane thinning is apparent around M protein. (d) Mean curvature of the membrane mapped onto the XY plane for outer (i) and inner (ii) leaflets showing small deformations around M from a nearly flat surface. (e) Accumulation of C1P and PI lipids in proximity to M_short_. (f) Residence time of C1P by residue in subunit A (above) and subunit B (below).

**Supplementary Figure 5.**
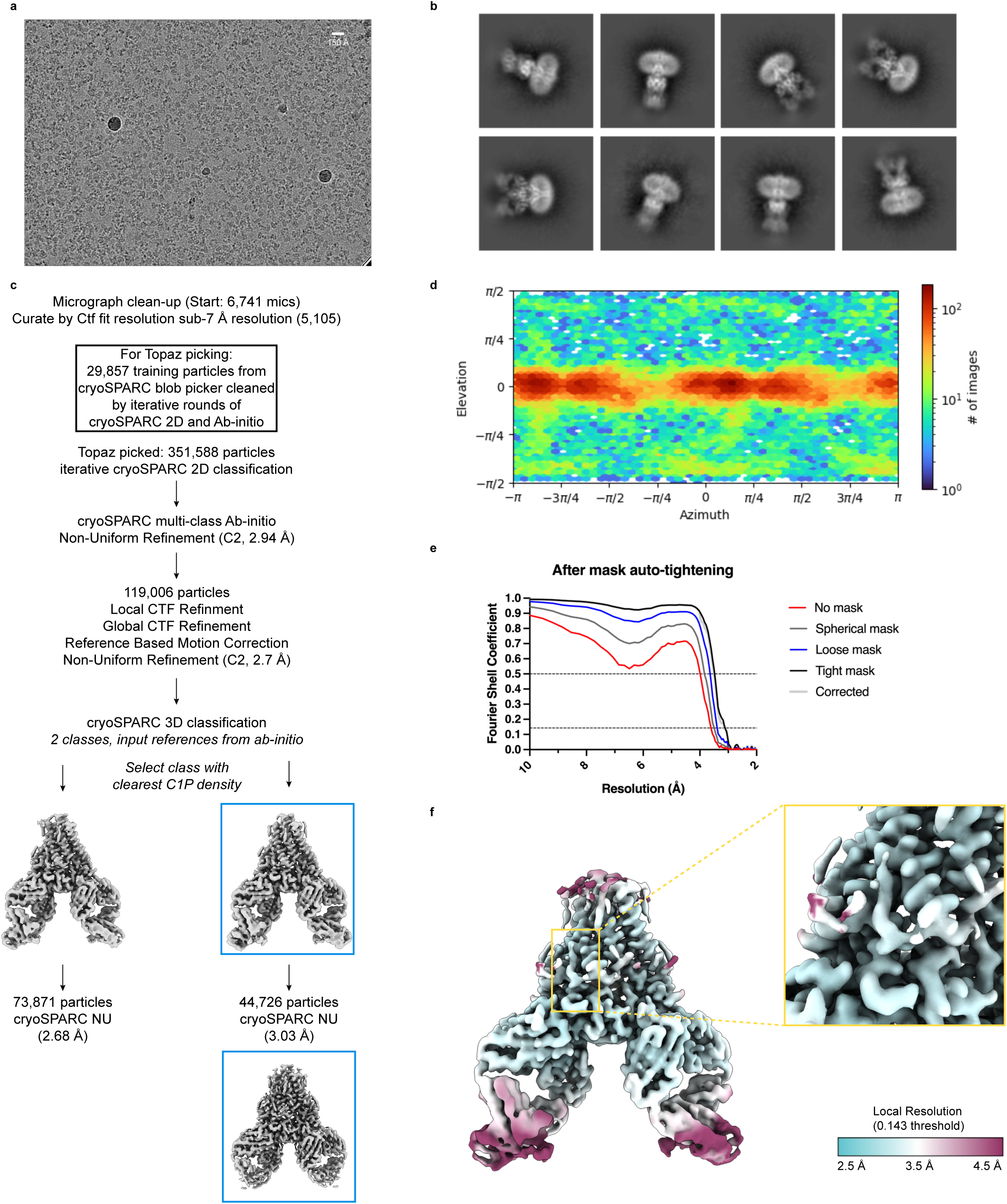
Cryo-EM data, processing pipeline, and validation for M_short_-C1P. (a) Representative micrograph, (b) selected 2Dclass averages, (c) cryo-EM data processing pipeline, (d) view angle distribution, (e) Fourier shell correlation (FSC) versus resolution between half maps from final non-uniform refinement, and (f) local resolution drawn on final sharpened map with inset zoomed in on C1P binding site.

**Supplementary Figure 6.**
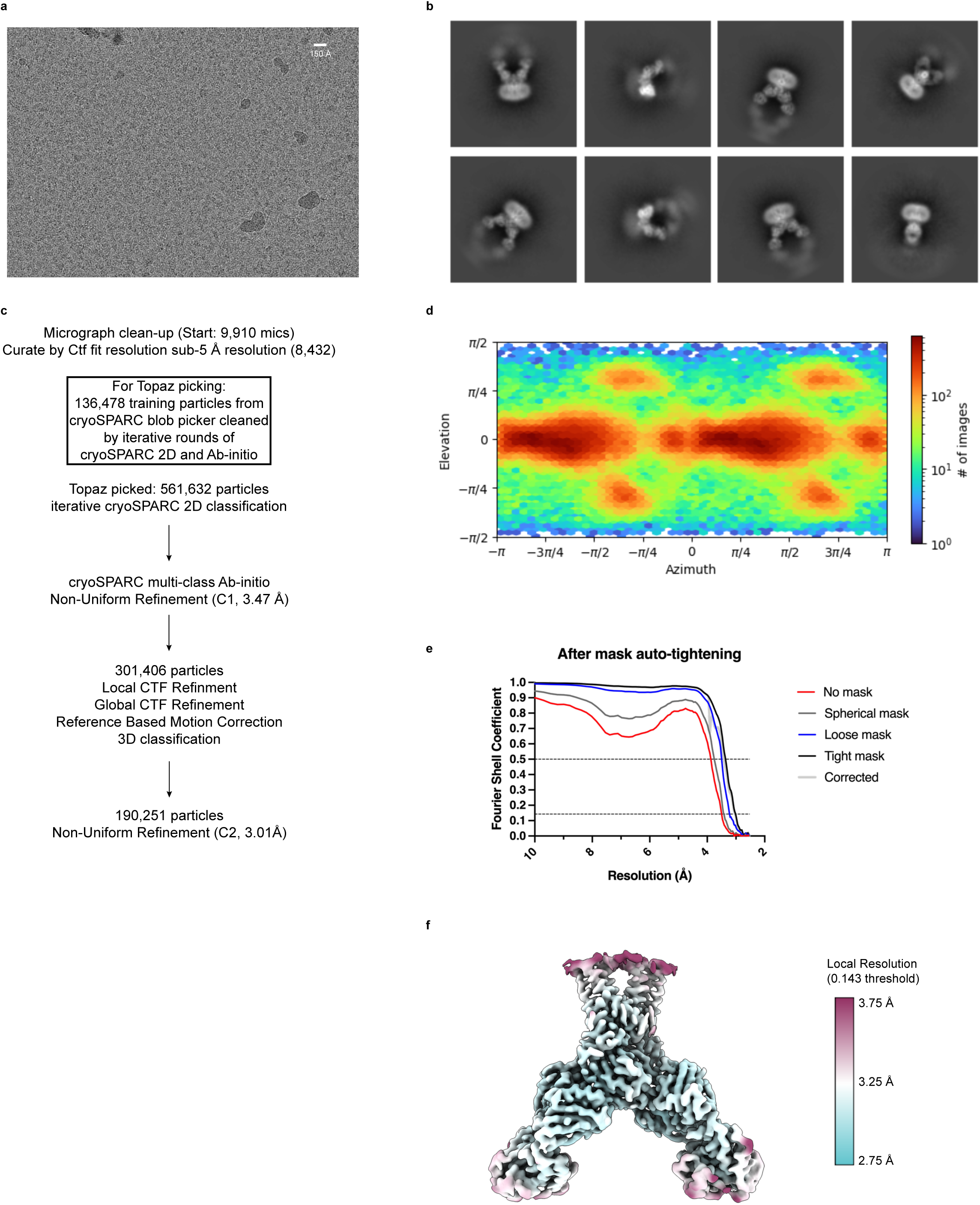
Cryo-EM data, processing pipeline, and validation for M_long_ in the presence of C1P. (a) Representative micrograph, (b) selected 2Dclass averages, (c) cryo-EM data processing pipeline, (d) view angle distribution, (e) Fourier shell correlation (FSC) versus resolution between half maps from final non-uniform refinement, and (f) local resolution drawn on final sharpened map.

**Supplementary Figure 7.**
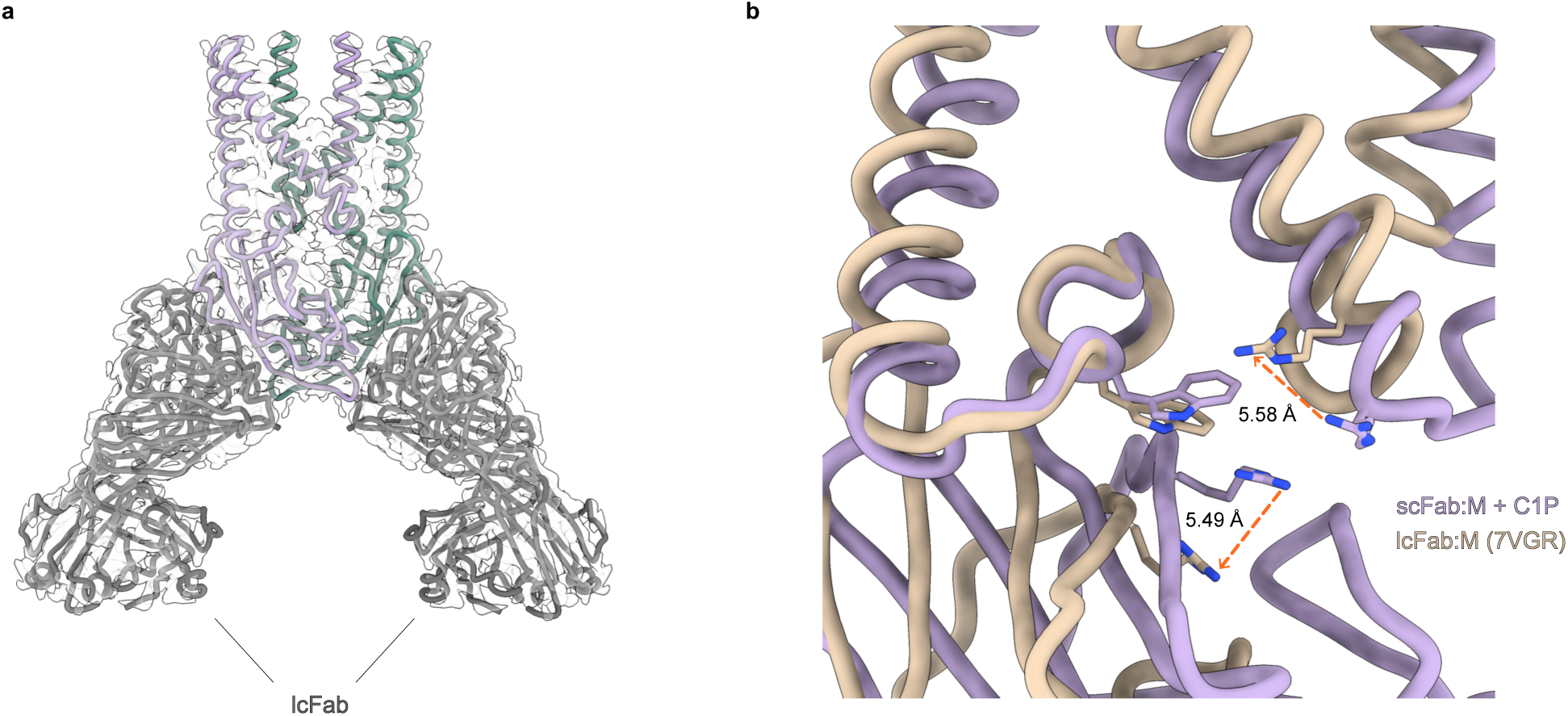
Structure of M_long_ in the presence of C1P. (a) Final model M in the long conformation viewed from the membrane. M subunits are purple and green and long conformation-selective Fabs are gray. 3.0 Å resolution sharpened map is shown as a transparent surface. (b) Overlay of M_long_ + C1P and M_short_-C1P structures highlighting rearrangement of the C1P binding site observed in M_short_ including relative displacement of key C1P-interacting residues.

**Supplementary Figure 8.**
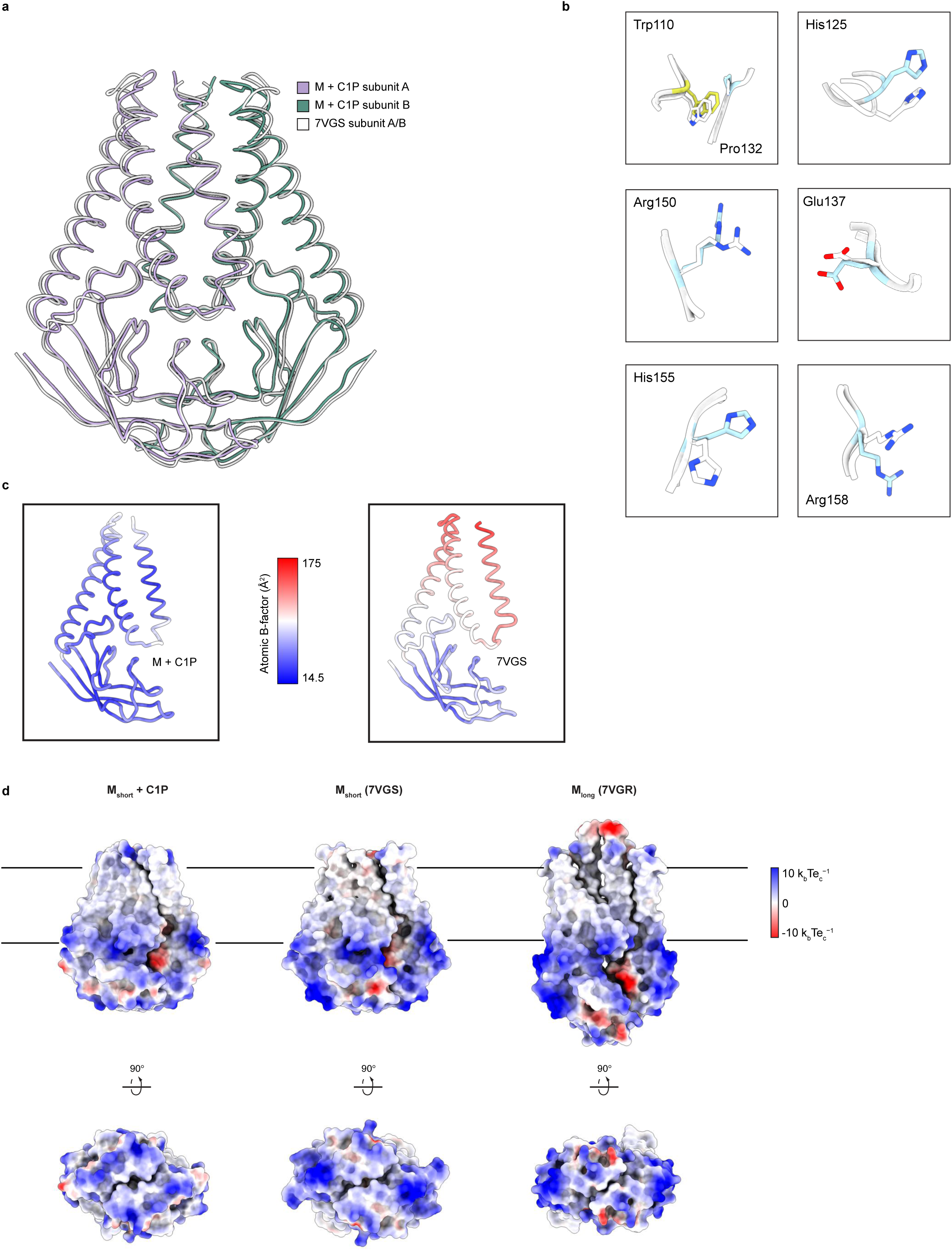
C1P binding compacts and stabilizes M_short_. (a) Overlay of M_short_+C1P (colored purple and green) and apo M_short_ illustrating compaction of transmembrane region. (b) Overlay of selected hinge and CTD residues from M_short_+C1P (blue) and apo M_short_ structures. (c) Single subunit from M_short_+C1P (left) and apo M_short_ (right) structures colored by refined B-factor. (d) M_short_+C1P (left), apo M_short_, and M_long_ electrostatic surface potential.

**Supplementary Figure 9.**
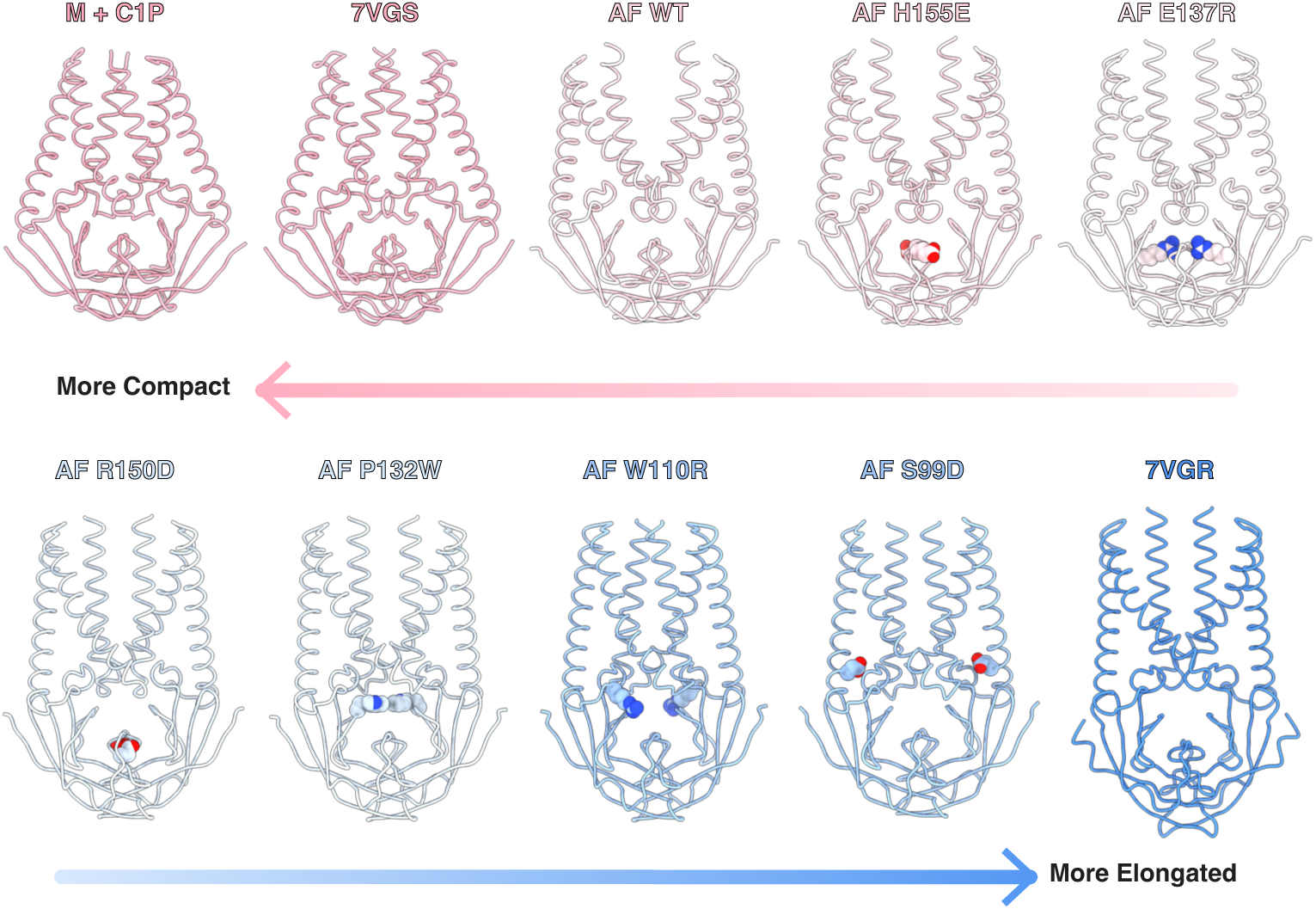
Mutations are predicted to shift M protein conformational landscape. Comparison of experimentally determined (M_short_-C1P, apo M_short_, and M_long_) and Aplhafold3-predicted (AF wild-type M, H155E, E137R, R150D, P132W, W110R, and S99D) structures. Mutated sites are drawn as spheres. Mutations were hypothesized to destabilize M_short_ based on M_short_-C1P and M_short_-JNJ-9676 structures. Models are arranged and colored from pink (most compact) to white (intermediate between short and long conformations) to blue (most elongated).

**Supplementary Figure 10.**
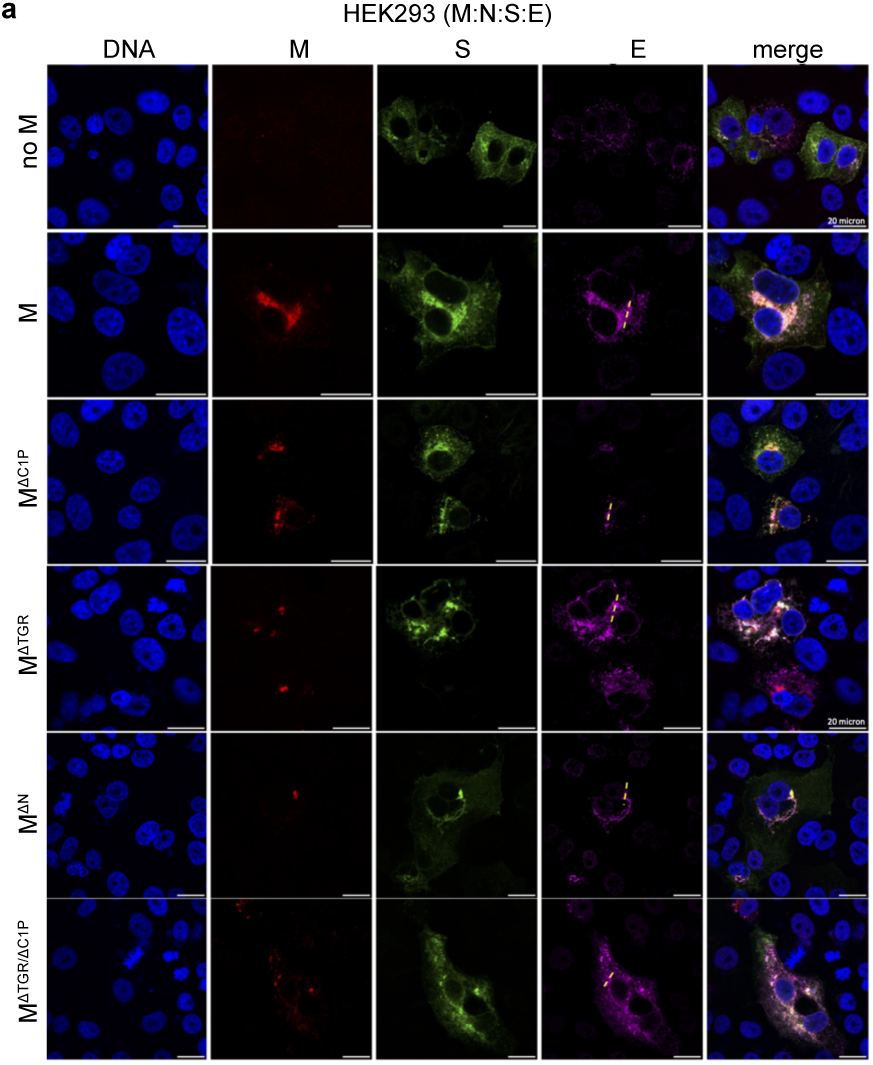
Colocalization of SARS-CoV-2 structural proteins in cells. HEK293 cells transfected with S,E,N, and M constructs used in this study and immunostained for M, S, and E.

**Supplementary Figure 11.**
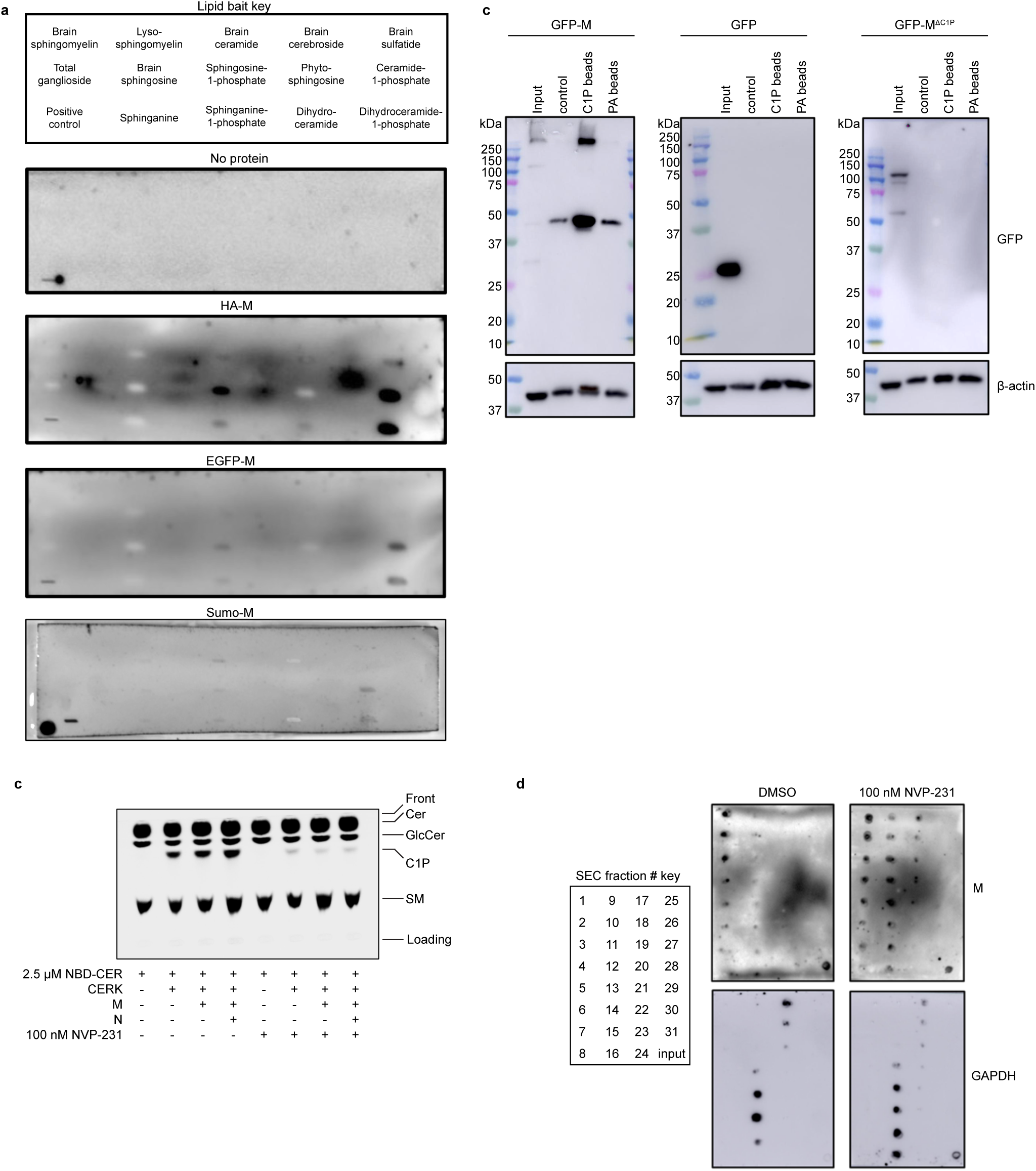
Uncropped blot and thin-layer chromatography images. (a) Sphingolipid snooper blot (Fig. 1A), (b) lipid-coated bead pull down blots (Figs. 1b, 4c), (c) CERK inhibition by NVP-231 assessed by TLC (Fig. 1e), and (d) blot of size chromatography fractionated lysates from DMSO- or NVP-231- treated cells expressing M, N, E, and S (Fig. 1e).

## Methods

### Molecular dynamics setup

The atomistic model of the M protein was built using the available cryo-EM structures of both the short (PDB ID: 7VGS) and long (PDB ID: 7VGR) conformations^19^. The CHARMM-GUI Membrane Builder ^53–56^ was employed to prepare initial MD systems, each consisting of a single M protein in either the short or long conformation embedded in a lipid bilayer. We utilized a 20×20 nm lipid bilayer membrane, which is sufficiently large to observe the membrane response of a single M protein. Based on biochemical experiments, we constructed two membrane models that mimic the composition of the endoplasmic reticulum-Golgi intermediate compartment ^23,24^. The first system is composed of a mixture of DYPC (45%), POPI (20%), POPE (15%), POPS (10%), and Cholesterol (10%). The second replaces POPS with C1P. We solvated the M/lipid-membrane system in a periodic rectangular box filled with TIP3P model water molecules, ensuring that the transmembrane region of the M protein was positioned close to the bilayer’s core. A multistep minimization and equilibration of the system was carried out following the protocols provided on CHARMM-GUI^57^. We used 0.15 M KCl to neutralize the system. We ran four conditions: M_short_ with C1P, M_short_ without C1P, M long with C1P, and M_long_ without C1P. Each condition was run in triplicate for twelve total MD simulations.

### Simulation details

All atomistic MD simulations were performed using GROMACS 2019 with CHARMM36m force field^58,59^. The simulations maintained a temperature of 310.15 K using a Nosé-Hoover thermostat ^60,61^ with a coupling time constant of 1.0 ps. During initial relaxation, the pressure was maintained at 1 Bar with a Berendsen barostat^62^. Periodic boundary conditions were applied with a time step of 2 fs. For the production run, the Parrinello-Rahman barostat was employed semi-isotropically due to the presence of a membrane^63,64^. A compressibility factor was set at 4.5×10^−5^ with a coupling time constant of 5.0 ps. Van der Waals interactions were computed using a switching function between 1.0 and 1.2 nm, and long-range electrostatics were calculated using the Particle Mesh Ewald (PME) method^65^. Hydrogen bonds were constrained using the LINCS algorithm^66^. We performed a 4 µs production run for each replica using Frontera (Texas Advanced Computing Center), and Midway2 (Research Computing Center at the University of Chicago) computing facilities.

### Free energy calculations

We used metadynamics to explore the free energy landscape^67^. The Hamiltonian of the system includes a history-dependent Gaussian bias potential, which is a function of a finite set of collective variables (CVs). In this study, we employed the well-tempered metadynamics (WT-MetaD) ^25^ technique to investigate the switching between the short and long conformational states of the M protein. In WT-MetaD simulations, the Gaussian bias height is progressively lowered to avoid overfilling the free energy basins and to achieve convergence ^68^.

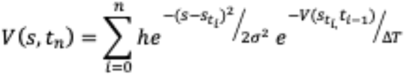

where h is the Gaussian height, σ is the Gaussian width, and *s_t__i_* are the sequence of points in CV space at time *t_t_*. ΔT controls the rate of height decrease.

From manual inspection of superimposed M_short_ and M_long_ structures, we defined two collective variables (CVs) to monitor conformational switching during simulations (Fig. 2a). The first reports the angle between CTDs and is calculated between S108 (subunit A), S184 (subunit A), and S108 (subunit B). This angle in M_short_ and M_long_ structures is 85° and 65°, respectively. The second reports on intersubunit separation and is defined as the distance between R107 in subunit A and subunit B. This distance in M_short_ and M_long_ structures 4.9 nm and 3.9 nm, respectively (Fig. 2a). We selected the Cα atom of each residue for angle and distance calculations.

In this work, Gaussians were deposited every 500 steps with an initial height set to 0.3 kJ mol^−1^. A bias factor of 15 was used to reduce the Gaussian height. The σ was set to 0.1. We conducted WT-metaD simulations using PLUMED 2.8.2 patched with GROMACS 2019 for both the model-C1P and control systems, using the long conformation as the starting structure. The error bars were computed using block averaging, as implemented in PLUMED.

### Analysis

Distances, angles and the number of contacts were calculated using the GROMACS module ^58^. Lipid distribution around the M protein and mean curvature were analyzed using the MDAnalysis Python package^69^. The residence time of interactions between lipid molecules and protein residues, as well as the occupancy of lipids around the protein, were determined using PyLipID^70^. Membrane thickness was calculated with VMD MEMBPLUGIN and Tcl scripts^71^.

### Protein cloning, expression, and purification

The coding sequence for SARS-Cov-2 M protein (Uniprot P0DTC5) was synthesized (IDT, Newark, NJ) and cloned into a vector based on the pACEBAC1 backbone (MultiBac; Geneva Biotech, Geneva, Switzerland) with an added C-terminal PreScission protease (PPX) cleavage site, linker sequence, superfolder GFP (sfGFP) and 7×His tag to generate the expression construct (SARS-CoV-2 M)-SNS-LEVLFQGP-SRGGSGAAAGSGSGS-sfGFP-GSS-7×His^18^. Bacmid was generated by transposition in transformed DH10MultiBac cells according to the manufacturer’s instructions.

Sf9 cells were cultured in ESF 921 medium (Expression Systems, Davis, CA) and P1 virus was generated from cells transfected with Escort IV reagent (MillaporeSigma, Burlington, MA) according to manufacturer’s instructions. P2 virus was generated by infecting cells at a density of 2 million cells/mL with P1 virus at a MOI ∼0.1. Infection was monitored by fluorescence and virus harvested at 72 hours. P3 virus was generated in a similar manner to expand the viral stock. The P2 or P3 viral stock was then used to infect Sf9 cells at 4 million cells/mL at a MOI ∼2–5. At 72 hours, infected cells containing expressed M-sfGFP protein were harvested by centrifugation at 2500 x *g* for 10 minutes and frozen at -80 °C.

Infected Sf9 cells from 1 L of culture (∼15 mL of cell pellet) were thawed in 100 mL of Lysis Buffer containing 50 mM HEPES pH 8.0, 150 mM KCl, 0.5 mM EDTA, 2 mM MgCl_2_, 2 mM DTT. Protease inhibitors (Final Concentrations: E64 (1 µM), pepstatin A (1 µg/mL), soy trypsin inhibitor (10 µg/mL), benzamidine (1 mM), aprotinin (1 µg/mL), leupeptin (1 µg/mL), AEBSF (1 mM), and PMSF (1 mM)) were added to the lysis buffer immediately before use. Benzonase (4 µl) was added after the cell pellet thawed. Cells were then lysed by sonication and centrifuged at 150,000 x *g* for 45 minutes. The supernatant was discarded, and residual nucleic acid was removed from the top of the membrane pellet using DPBS. Membrane pellets were resuspended in High Salt Lysis buffer (50 mM HEPES pH 8.0, 1 M KCl, 1 mM EDTA, 2 mM DTT) plus protease inhibitors, followed by ultracentrifugation at 150,000 x *g* for 45 minutes. The supernatant was discarded, membrane pellets were washed again with DPBS, then transferred into a Dounce homogenizer containing extraction buffer (50 mM HEPES, 150 mM KCl, 1 mM EDTA, 1% lauryl maltose neopentyl glycol (LMNG, Anatrace, Maumee, OH), 0.1% cholesteryl hemisuccinate Tris salt (CHS, Anatrace, Maumee, OH) pH 8). A stock solution of 10% LMNG, 1% CHS was dissolved and clarified by bath sonication in 200 mM HEPES pH 8 prior to addition to buffer to the indicated final concentration. Membrane pellets were homogenized in extraction buffer and this mixture (150 mL final volume) was gently stirred at 4°C for 1.5 hours. The extraction mixture was centrifuged at 33,000 x *g* for 45 minutes and the supernatant, containing solubilized membrane protein, was bound to 7.5 mL of Sepharose resin coupled to anti-GFP nanobody for 1.5 hours at 4°C. The resin was then collected in a column and washed with 100 mL of buffer 1 (20 mM HEPES pH 8.0, 150 mM KCl, 1 mM EDTA, 0.005% LMNG, 0.0005% CHS), 150 mL of buffer 2 (20 mM HEPES pH 8.0, 500 mM KCl, 1 mM EDTA, 0.005% LMNG, 0.0005% CHS), and 25 mL of buffer 1. The resin was then resuspended in 7.5 mL of buffer 1 with 0.5 mg of PPX protease and rocked gently in a capped column for 2 hours. M protein was then eluted with an additional 15 mL of buffer 1, spin concentrated to ∼1 mL (Amicon Ultra 30 kDa cutoff, Millipore, Tullagreen, Ireland), and loaded onto a Superose 6 increase column (GE Healthcare, Chicago, IL) on an NGC system (Bio-Rad, Hercules, CA) equilibrated in buffer 1. Peak fractions containing M protein were stored overnight at 4 °C prior to preparation of Fab complexes.

Coding sequences for conformationally selective antibody fragments (lcFab and scFab)^19^ were synthesized and cloned in a pTwist CMV BetaGlobin WPRE Neo vector with C-terminal 7X-His tags on heavy chain (Twist Biosciences). Fabs were cotransfected in suspension HEK293T GNTI-cells for expression using PEI according to manufacturer’s instructions. Supernatants were thawed on ice, buffered with HEPES pH 8.0 at a final concentration of 100 mM, and centrifuged at 21,000 x *g* for 10 minutes to pellet any precipitated material. Clarified supernatant was incubated with 3 mL His-Pure Ni NTA resin (ThermoFisher Scientific, Rockford, IL) for 1.5 hours at 4°C. The Ni NTA resin was collected in a column and washed with 30 mL each of Cholate buffer (40 mM HEPES pH 8.0, 300 mM KCl, 50 mM Na-cholate, 10 mM Imidazole) and Imidazole Wash Buffer (40 mM HEPES pH 8.0, 300 mM KCl, 20 mM Imidazole). The Fabs were eluted in 4 column volumes of Ni NTA elution buffer (40 mM HEPES pH 8.0, 300 mM KCl, 200 mM Imidazole). The eluate was spin concentrated to ∼1 mL with Amicon Ultra spin concentrator 30 kDa cutoff (Millipore, Tullagreen, Ireland) and loaded onto a Superdex 200 increase (S200) column (GE Healthcare, Chicago, IL) on an NGC system (Bio-Rad, Hercules, CA) equilibrated in SEC buffer (20 mM HEPES pH 8.0, 150 mM KCl, 1 mM EDTA). Fractions containing complete Fabs were pooled, concentrated, and stored at 4°C for future use.

### M-Fab complex formation and purification

The peak fraction from size exclusion chromatography containing pure homodimeric M protein was collected, incubated with purified lcFab or scFab at a 1:2.2 M monomer:Fab molar ratio for 45 minutes on ice, and run on a Superose 6 increase column in Buffer 1 to separate excess Fabs. Pooled peak fractions containing M:Fab complexes were spin concentrated with an Amicon Ultra 50 kDa cutoff spin concentrator to 2.2 mg/mL (scFab) and 2.7 mg/mL (lcFab). Concentrated complex samples were cleared by a 10-minute 21,000 x g spin at 4 °C, then d18:1/16:0 ceramide-1-phoshate (C1P; N-palmitoyl-ceramide-1-phosphate ammonium salt, (Avanti Polar Lipids, Alabaster, AL)) was added to each sample to a final concentration of 100 µM from a 1 mM stock in 1:1 MeOH:CHCl_3_. Samples were then incubated on ice for 30 minutes prior to grid preparation.

### Cryo-EM sample preparation and data collection

3.4 µl samples of M:lcFab or M:scFab in LMNG/CHS and C1P were applied to freshly glow discharged Holey Carbon, 300 mesh R 1.2/1.3 gold grids (Quantifoil, Großlöbichau, Germany) (M:scFab) or C-Flat Au 300 mesh 1.2/1.3 (Electron Microscopy Sciences, Hatfield, PA) (M:lcFab) then plunge frozen in liquid ethane using a FEI Vitrobot Mark IV (ThermoFisher Scientific) with a ∼5 second wait time set to 4°C, 100% humidity, blot force 1, and 3 second blot time.

Grids were clipped and cryo-EM micrographs were collected on a Titan Krios G3i microscope operated at 300 kV. Fifty frame videos with a total electron dose of 50 e-/Å^2^ were recorded on a Gatan K3 Summit direct electron detector. The M:scFab dataset was collected in super resolution mode with a pixel size of 0.4055 Å (0.8110 Å physical pixel size), and the M:lcFab dataset was collected in counting mode with a pixel size of 0.848 Å. Videos were recorded around a central hole position via image shift in an 11×11 hole pattern with two targets per hole. Target defocus range was -0.5 to -1.8 µm. See Supplemental Table 4.1 for detailed data collection statistics.

### Cryo-EM data processing

Data processing for both samples were done in cryoSPARC v4.4.1^72^. For M:scFab in LMNG/CHS/C1P 6,741 movie stacks were collected, motion-corrected, binned to 0.8110 Å pixel^-1^ using Patch Motion Correction, and CTF-corrected using Patch CTF Estimation (Supplementary Fig. 5). Micrographs with an estimated CTF resolution below 7 Å were discarded, leaving 5,105 for further processing. Initial particle picking was performed using cryoSPARC’s reference free blob picker and curated from 404,953 particles down to 29,857 ‘good’ particles by 2D classification.

Topaz training, picking and extraction yielded 351,588 particles which were then iteratively 2D classified, resulting in a set of 163,911 ‘good’ particles. A two-class ab-initio reconstruction was performed to further sort out 44,905 particles that did not contribute to a high-resolution 3D map. Subsequent non-uniform refinement (C1, 2 extra passes, 14 Å initial resolution) produced a map with an overall 2.94 Å resolution showing clear lipid densities on both leaflets. This map and corresponding 119,006 particles were subjected to local CTF refinement, then global CTF refinement, followed by polishing via Reference Based Motion Correction which further improved the resolution to 2.7 Å with improved lipid densities. A round of 3D classification (2 classes, 3 Å target resolution, 10 O-EM epochs, 0.95 class similarity) and subsequent C2 non-uniform refinement on the class with the clearest lipid densities produced a final 3.03 Å resolution map (44,726 particles) where a C1P molecule could be unambiguously modeled (Supplementary Fig. 5, Supplementary Table 1). The same non-uniform refinement with C1 symmetry produced a 3.26 Å resolution map with no apparent differences between subunits or lipid binding, so the C2 map was chosen for building an atomic model as the C1P densities were more complete and clearly defined.

The M:lcFab LMNG/CHS/C1P dataset was processed in a similar manner, with 9,910 initial micrographs culled to 8,432 using a 5 Å cutoff. Topaz was trained on 136,478 particles yielding a stack of 561,632 particles, which was cleaned, refined and polished as described for the scFab dataset, yielding 301,406 ‘shiny’ particles. A round of 3D classification yielded a class of 190,251 particles with 3.01 Å final resolution after non-uniform refinement with C2 symmetry (Supplementary Fig. 6, Supplementary Table 1). Further attempts at 3D classification and heterogenous refinement failed to separate these particles into subsets with unique characteristics or improved map quality.

### Modeling, refinement, and analysis

Modeling of the M:scFab complex in LMNG/CHS/C1P was performed in Coot ^73^ using cryoSPARC sharpened cryo-EM maps, the SARS-CoV-2 M protein short conformation structure (PDB ID: 8CTK), and an AlphaFold3^74^ model of the scFab variable domains rigid body fitted into the map in Coot as initial models. The C1P ligands (3 letter code 1PX) were added in Coot. The model was real space refined in Phenix ^75^ and validated using Molprobity^76^. Mutant models were generated with AlphaFold3.

We assessed putative metal coordination using several computational tools. First, AlphaFold3^74^ structure predictions of M dimers with two cations (potassium, sodium, calcium, and magnesium) all show ion occupancy of the putative binding site with minimal distortions to the overall fold. Second, Metric Ion Classification (a deep learning tool for assigning ions in cryo-EM maps^77^) and CheckMyMetal (CMM, an ion validation tool that accounts for ligand chemistry, geometry, and refined B-factor statistics^78^) both predict metal over water binding to this site. We therefore separately modeled each ion in the structure, refined the model in Phenix ^75^ and compared validation statistics in CMM. The site was predicted in all cases to be most likely occupied by potassium and K+ is included in the final models.

### Lipid binding assays

HEK293 cells were used to express and purify M protein for lipid binding overlay assays. Eight µg of DNA was diluted into Opti-MEM, mixed with PEI in the ratio of 3:1 (PEI:DNA), and incubated for 10 minutes at room temperature. The PEI-DNA mixture was added to cells plated in a 60 mm dish at 80% confluency. 5 hours post-transfection the media was changed to DMEM + 10 % FBS. The cells were then incubated for 24 hours at 37°C and 5% CO_2_. Media was then removed from the dish and cells were washed twice with PBS and lysed with lysis buffer (10 mM HEPES pH 7.4, 150 mM NaCl, 0.25% Igepal CA-630, 5 µM PMSF, 5 mM EDTA, and Halt protease inhibitor cocktail). Cells were incubated in lysis buffer at 4°C for 10 min. Cells were then scraped from the dish and transferred to a prechilled centrifuge tube. The cell lysate was incubated for 30 min at 4°C with frequent 30-40 sec vortexing. The cell lysate was then centrifuged at 16,000 x *g* for 10 min at 4°C to separate the soluble and insoluble fractions. The supernatant was transferred to a microcentrifuge tube and small aliquots were used to determine the protein concentration with the BCA method. SDS-PAGE followed by Western blot was performed on the purified cell lysates to confirm M protein expression and check expression levels for different conditions/replicates. An anti-M antibody was used for M protein detection and an anti-actin antibody was used for control blots on cell lysates.

Lipid strips or sphingolipid snoopers were used to detect binding of M protein to different lipids. 0.5 µg of cell lysate was added as a control to one corner of each membrane to ensure M detection in each blot. The membrane was allowed to fully dry after cell lysate blotting. Once dry, the membrane was saturated with 1% milk in TBS-T for 1 hr at room temperature. After 1 hour, the solution was replaced with 4 µg/ml of total cell lysate in 1% milk in TBS-T and incubated overnight at 4°C. The membrane was then washed three times for 10 min each with TBS-T at room temperature. The membrane was incubated with anti-SARS-COV-2 M protein antibody diluted 1:5000 in 1% milk in TBS-T for 1 hr at room temperature. The membrane was then washed three times for 10 min each with TBS-T at room temperature. The membrane was then incubated for 1 hour at room temperature with secondary antibody (goat anti-rabbit) diluted at 1:5000 in 1% milk in TBS-T for 1 hr at room temperature. M protein interactions with different lipid classes was resolved using ECL Clarity reagents and an imager (Amersham Imager 600) to visualize M protein location on each membrane.

To determine the ability of M protein to interact with lipid coated beads, a pull down assay was performed using control beads, C1P beads, and PA beads (Echelon Biosciences, Inc., Salt Lake City, UT) and adapted from a published lipid coated bead binding protocol (https://bio-protocol.org/en/bpdetail?id=1039&type=0). M protein was expressed in HEK293 cells for 24 hours and cells were collected and lysed, which was followed by low speed centrifugation to remove cellular debris. Beads were incubated with cell lysates for 30 minutes at room temperature and the lipid bound and unbound fractions were separated centrifugation. SDS-PAGE analysis was performed on the bead pull down samples, which was followed by Western blot to detect the M protein in each sample.

### C1P cellular assays

TLC plates were prepared with 20 cm height and width and incubated with 50 of 1:1 chloroform:methanol (v/v) in a glass tank. The TLC plate was secured in a plate holder for stability and the tank lid was closed until the solution ascended to 0.5 cm from the top of the plate (usually 2 to 3 hrs depending on room temperature). The TLC plate was allowed to dry for 5 min in a fume hood. The TLC plate was then activated for 30 minutes in an oven at 100°C. Lipid samples were prepared in chloroform (5-10 µl), sonicated, spun down, and loaded (0.5 to 2 µl of each sample) ∼1 cm from the bottom of the TLC plate. Spots were dried prior to loading the plate in a saturated tank. Plates were sprayed with primuline (0.005% (w/v) in 4:1 acetone:water (v/v)), dried in the fume hood, and imaged at 345 nm (Amersham Imager 600). To assess M protein chromatography profiles with and without inhibition of CERK, cells were treated with vehicle (DMSO) or 100 nM NVP-231 and lysed and for analysis of M protein size using a Superdex^TM^ 200 Increase small-scale SEC column (Cytiva Life Sciences, Marlborough, MA). Fractions were collected every 1 mL and a portion of each fraction was spotted onto nitrocellulose to determine M protein content in each fraction via detection of M using Western blot.

### Cellular imaging and fractionation assays

HEK293, COS-7, or Vero E6 cells were used for SARS-CoV-2 protein expression and detection with confocal microscopy. Transfections were performed for M, N, S, and Flag-E alone or in combinations to assess M protein localization and colocalization with S or E. Imaging analysis was performed 16-24 hours post-transfection. Images were analyzed in Image J for localization and co-localization analysis. The cellular synctia assay was adapted from Zhang et al. ^79^. 24 hours post-transfection of S, S and M, or S with different M mutations, were imaged post nuclei staining and percent synctia formation was determined as the percentage of nuclei with fused cells.

### VLP budding and entry assays

VLP production was performed and modified from Plescia et al. ^17^. Transfections of plasmid DNAs of M:N:S:Flag-E at a 6:3:8:8 ratio in 50 μl Opti-MEM, mixed with PEI in a 3:1 ratio (PEI:DNA), and incubated for 10-15 min at room temperature, and added to cells. Cells were placed at 37°C and 5% CO_2_ and five to six hours post-transfection, the media was changed to DMEM + 10 % FBS. Transfected cells were incubated at 37°C and 5% CO_2_. 48 hours post transfection, supernatants were collected in a chilled falcon tube with 1× Halt^TM^ protease inhibitor cocktail. Cells were washed with 1× PBS and centrifuged 10 min 4° C at 1000 x *g*. The supernatant was transferred to a new falcon and centrifuged 10 min, 4° C at 1000 x *g* and filtered through 0.45 micron filter into a new tube. Supernatants were deposited on a 20% sucrose cushion.

VLPs were centrifuged at 28,000 rpm for 3 hours at 4°C. 4. The resulting VLPs were used for entry assays and quantified for protein content using a BCA assay. VLP entry assays were performed using equal amounts of VLPs for each condition. Cells subjected to VLPs were spinoculated for 45 min at 4°C, and allowed to incubate for 1 hour at 37°C. Plates were then rinsed with PBS and nuclei were stained with Hoechst 33342 for 15 min at 37 °C prior to fixation with 4% paraformaldehyde (PFA) and confocal imaging. Quantification of VLP entry signal was performed by calculating the total number of VLPs (green puncta)/the total number of cells.

Figures were prepared using VMD, PyMOL, ChimeraX^80^, Prism, Python, Matlab, and Adobe Illustrator software.

## Acknowledgements

This work was funded by the National Institutes of Allergy and Infectious Diseases grant R01-AI169896. Computational resources were provided by the Frontera Supercomputer at the Texas Advanced Computer Center (TACC) and the Research Computing Center (RCC) at the University of Chicago.

## Author contributions

M.D. performed and analyzed molecular dynamics simulations with G.A.V. K.A.D. performed and analyzed cryo-EM and structure prediction experiments with S.G.B. S.A., E.J.B., and R.S. performed and analyzed lipid binding, microscopy, and virus-like particle experiments with R.V.S. S.G.B., R.V.S., G.A.V., K.A.D., and M.D. wrote the paper with input from all authors.

